# TGF-β serves as a critical signaling determinant of liver progenitor cell fate and function

**DOI:** 10.1101/2025.06.04.657336

**Authors:** Chenhao Tong, Tao Lin, Han Wang, Carolina De La Torre, Hui Liu, Chen Shao, Seddik Hammad, Roman Liebe, Matthias P Ebert, Huiguo Ding, Steven Dooley, Hong-Lei Weng

**Affiliations:** Department of Medicine II, Section Molecular Hepatology, University Medical Center Mannheim, Medical Faculty Mannheim, Heidelberg University, Mannheim, Germany; Hepatopancreatobiliary Surgery Department, The First Affiliated Hospital of Ningbo University, Ningbo, Zhejiang, China; Department of Biochemistry, Case Western Reserve University, Cleveland, OH, USA; NGS Core Facility, Medical Faculty Mannheim, University of Heidelberg, Mannheim, Germany; Department of Pathology, Beijing You’an Hospital, Affiliated with Capital Medical University, Beijing, China; Clinic of Gastroenterology, Hepatology and Transplantation Medicine, Essen, Germany; Department of Medicine II, University Medical Center Mannheim, Medical Faculty Mannheim, Heidelberg University, 68167-Mannheim, Germany; DKFZ-Hector Cancer Institute at the University Medical Center, Mannheim, Germany; Molecular Medicine Partnership Unit, European Molecular Biology Laboratory, Heidelberg, Germany; Department of Gastroenterology and Hepatology, Beijing You’an Hospital, Affiliated with Capital Medical University, Beijing, China

**Author notes:** **Corresponding author:** Hong-Lei Weng Department of Medicine II, Section Molecular Hepatology, University Medical Center Mannheim, Medical Faculty Mannheim, Heidelberg University. Theodor-Kutzer Ufer 1-3, 68167 Mannheim, Germany.

**Keywords:** HGF, EGF, SMAD, Massive hepatic necrosis, Acute liver failure

## Abstract

Liver progenitor cells (LPCs) are the smallest cholangiocytes that perform hepatocyte functions to rescue the lives of patients suffering from acute liver failure (ALF) caused by massive hepatic necrosis (MHN). To date, it remains largely unknown how LPCs remain quiescent and become activated following MHN. This study elucidates the essential role of TGF-β in regulating LPC quiescence and activation. Spatial transcriptomics analysis of liver tissues from four MHN-ALF patients revealed that LPCs receive multiple active signals from surrounding macrophages and hepatic stellate cells, including TGF-β, HGF, and EGF. Physiologically, TGF-β inhibits LPC proliferation by impeding G1-S phase transition. Ectopic Smad7 expression remarkably increased LPC proliferation in 3,5-diethoxycarbonyl-1,4-dihydrocollidine-fed mice. Intriguingly, extensive LPC proliferation was observed in ALF patients despite robust TGF-β-p-SMAD2 signaling in activated LPCs. Immunohistochemistry and immunofluorescence revealed significantly elevated expression of p-MET, p-STAT3, p-EGFR, and p-ERK in LPCs, indicating active HGF and EGF signaling. In vitro, either HGF or EGF promoted LPC proliferation despite the presence of TGF-β. Beyond acting as mitogens, HGF and EGF regulate master hepatocyte genes (e.g., HNF4α) and cholangiocyte genes (e.g., SOX9) in LPCs. Notably, HGF-dependent HNF4α required TGF-β-activated SMADs. Collectively, TGF-β serves as a critical signaling determinant of LPC fate and function.

---

Liver progenitor cells (LPCs), the smallest cholangiocytes, constitute a liver specific stem-cell like population that performs vital liver functions to support systemic homeostasis in acute liver failure (ALF) following massive hepatic necrosis (MHN), the most severe pathologic lesion in ALF^1-5^. The performance of these cells determines whether an ALF patient can survive the golden window - the first 7 days since the initiation of disease^5^. Over time, the activated LPCs differentiate into mature hepatocytes and restore lost liver mass for the organ recovery^5^. Therefore, LPCs mediate the second pathway of liver regeneration in the condition of massive hepatocyte loss^6^. To date, the detailed mechanisms of LPC biology, including how LPCs rapidly proliferate, perform hepatic function, and subsequently differentiate into hepatocytes, are largely unknown.

Physiologically, quiescent LPCs reside in the Canal of Hering and the smallest biliary tree^1^. When they are stimulated by inflammatory mediators in liver disease, activated LPCs form ductular reaction (DR)^1^. In different disease settings, DR may arise from different cell sources, including proliferation of pre-existing cholangiocytes, LPCs, and rarely, biliary metaplasia of hepatocytes^7^. In ALF associated with MHN, LPCs are the predominant cell source of DR because most parenchymal cells are destroyed in such an urgent clinical syndrome. There are three essentially open questions in LPC biology: (1) How are LPCs kept in a quiescent state in the normal liver? (2) Which microenvironment-derived external signals such as inflammatory mediators induce LPC proliferation? (3) How do these external signals drive LPC differentiation towards hepatocyte? To date, multiple inflammatory cytokines, including IL-6, TNF-α/TWEAK, IFN-γ and TGF-β, have been reported to regulate LPC activation in different liver disease and animal models^8-12^. Among these factors, the effects of TGF-β on LPC biology are worth further in-deep investigation.

TGF-β superfamily members, TGF-β, activin and bone morphogenic protein (BMP), play fundamental roles in multiple basic biological processes of humans, including development, tissues homeostasis and regeneration^13^. TGF-β signaling is regarded as a guardian in the preservation of systemic, tissue and cell integrity^14^. To date, the effects of TGF-β on cell proliferation have been investigated in multiple cells. In many types of normal cells, TGF-β has the last word in determining whether these cells proliferate^15^. In the context of cell cycle control, the most important targets of action by TGF-β are the genes CDKN1A and CDKN2B, which encode the two CDK inhibitors p15^INK4B^ and p21^Cip1^, respectively^16,17^. In this setting, transcription factors FOXO1/3/4, C/EBPβ and Miz-1 are required for the SMAD3-SMAD4 complexes to activate transcription of p15^INK4B^ and p21^Cip1^ ^16,17^. In addition, that C/EBPβ synergizes SMAD proteins to repress c-MYC also contributes to inhibit cell proliferation^17^. In normal rat livers, inhibition of TGF-β receptors by dominant negative plasmids results in hepatocyte proliferation^18^, suggesting an anti-proliferative effect of TGF-β on liver cells. Besides regulating liver cell proliferation, TGF-β also regulates multiple liver physiological and pathophysiological processes. In liver development, TGF-β provides a crucial signaling, which is essential for hepatoblast differentiation towards intrahepatic cholangiocytes^19^. In adult livers, TGF-β drives the formation of novel biliary system from hepatocytes in a mouse model mimicking Alagille Syndrome^20^. In addition to regulate cholangiocytes, TGF-β signaling contributes to hepatocyte function. Transcription of master hepatocytic transcription factor hepatocyte nuclear factor 4α (HNF4α) in hepatocytes requires TGF-β to provide activated SMADs, which bind to the gene promoter and recruit the histone acetyl-transferase CREB-binding protein (CBP)/p300^21^. The TGF-β-SMAD signaling is detectable in the majority of liver cell types, including hepatocytes, cholangiocytes, macrophages and HSCs^22^. The crucial role of TGF-β in liver fibrosis has been well recognized for decades^23-25^. However, the exact mechanisms regarding how the TGF-β-SMAD signaling regulates LPCs in ALF following MHN remains largely unknown to date.

In this study, we performed a spatial transcriptomics analysis on liver tissues collected from four irreversible ALF patients undergoing liver transplantation. Cell-cell interaction analyses revealed predominant signaling pathways in LPCs (e.g., TGF-β, HGF, and EGF), which are produced by macrophages, activated hepatic stellate cells (HSCs) or both. In normal LPCs, TGF-β inhibited LPC proliferation through impeding G1-S phase transition, while in ALF, HGF and EGF prevails TGF-β-dependent proliferative inhibition, thereby leading to LPC proliferation. Although the anti-proliferative effect of the TGF-β-SMAD signaling was compromised, activated SMADs were essential for LPCs to initiate the transcription of vital hepatocytic genes, including HNF4α.

## Results

### Macrophages and activated hepatic stellate cells are predominant in the ductular reaction

We first conducted histological examination, including hematoxylin & eosin (HE), immunohistochemistry (IHC) and immunofluorescent (IF) staining, in 8 irreversible ALF patients who underwent liver transplantation (**Figure 1A**). DRs were characterized by proliferative LPCs surrounding by remarkable inflammatory cell infiltration (**Figure 1B**). IHC for CD68 and α-SMA demonstrated that LPCs were surrounded by numerous macrophages and activated hepatic stellate cells (HSCs) (**Figure 1C**). As LPCs differentiated into intermediate hepatocyte-like cells (IHLCs) and mature hepatocytes (**Figure 1D**), macrophages and activated HSCs gradually decreased (**Figure 1E**). These results suggested that macrophages and activated HSCs may play critical roles in LPC proliferation and differentiation.

**Figure 1.**
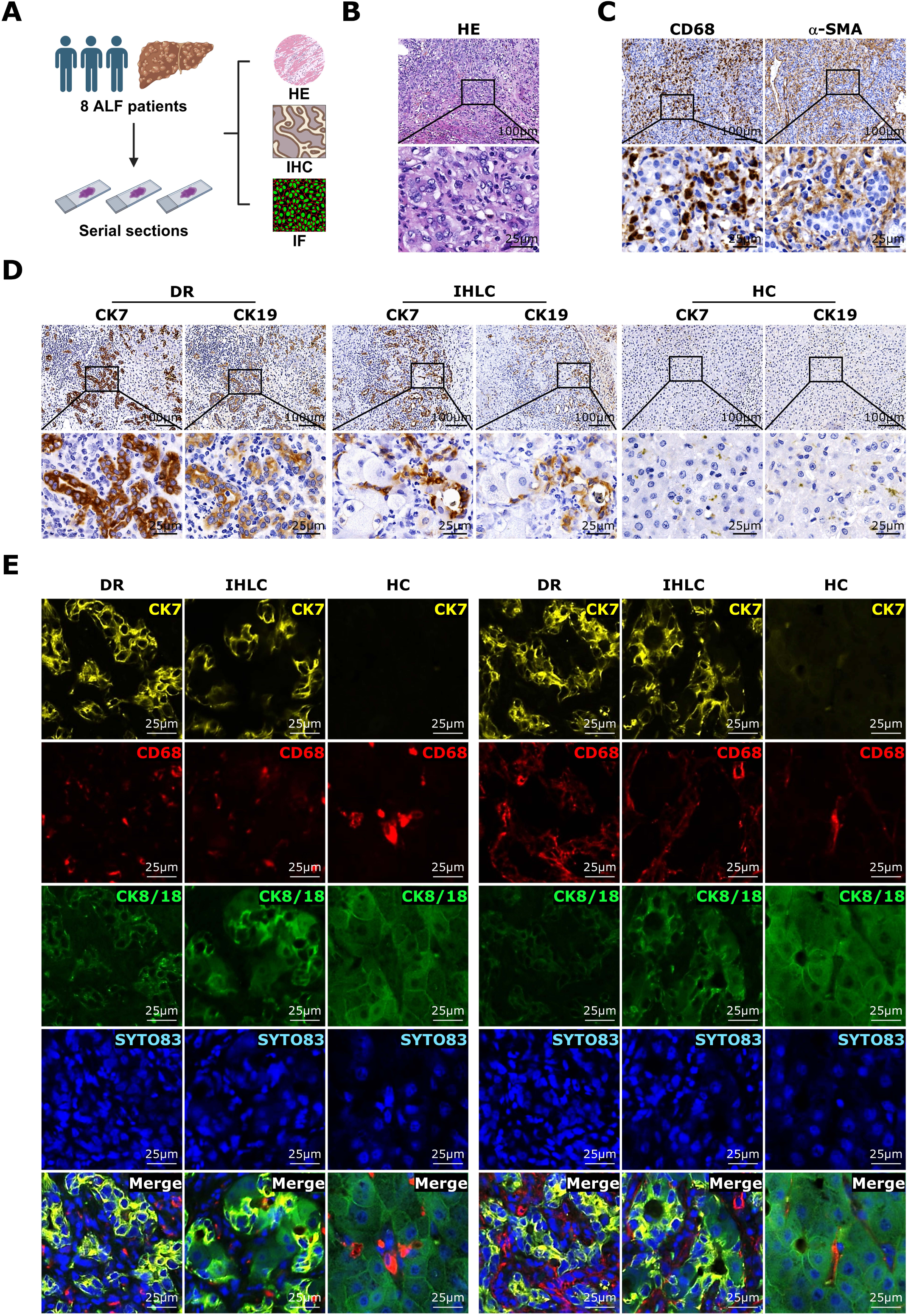
Macrophages and activated hepatic stellate cells are predominant in the ductular reaction of acute liver failure patients. **(A)** Hematoxylin and eosin (HE), immunohistochemistry (IHC), and immunofluorescence (IF) were performed on liver tissues collected from 8 irreversible acute liver failure (ALF) patients who underwent liver transplantation. **(B)** A representative HE staining image shows an area with massive hepatocyte loss. A frame highlights ductular reaction (DR) regions, where proliferative liver progenitor cells (LPCs) are surrounded by dense inflammatory cell infiltration. **(C)** IHC staining for CD68 and α-SMA was performed to highlight macrophages (Mø) and activated HSCs in DR regions. **(D)** IHC staining for CK7 and CK19 was performed to demonstrate the transition from DR and intermediate hepatocyte-like cells (IHLCs) to mature hepatocytes (HCs). **(E)** Co-IF staining for CK7 (yellow), CD68 or α-SMA (red), CK8/18 (green), and nuclei (SYTO83, blue) was used to identify LPCs, macrophages, activated hepatic stellate cells (HSCs), HCs hepatocytes in ALF patients. Scale bars: 100 μm (B–D, low magnification); 25 μm (B–E, high magnification).

### Spatial transcriptomic analysis reveals transcriptome of liver progenitor cells, macrophages and hepatic stellate cells in ALF patients

Next, we performed a GeoMx digital spatial transcriptomics analysis on liver tissues collected from 4 irreversible ALF patients (**Figure 2A**). We selected multiple regions of interest (ROIs) from different cell types, including CK7-positive LPCs (n=37), CK8/18-positive hepatocytes (n=20), CD68-positive macrophages (n=37), and α-SMA-positive HSCs (n=21) **(Figure 2A)**. Principal component analyses (PCA) based on the chosen ROIs revealed completely distinct transcriptomic profile between LPCs and HCs (**Figure 2B**). There were partially overlapping transcriptomic profiles between macrophages and activated HSCs (**Figure 2B**). LPCs exhibited high levels of EPCAM, SOX9, KRT7 and KRT19 transcripts, whereas hepatocytes expressed CYP2E1, albumin, HNF1A, HNF4A and CEBPA transcripts (**Figure 2C**). Macrophages showed high levels of CD68, CD14, CD86, and CD163 transcripts, whereas activated HSCs expressed ACTA2, MMP2, PDGFRB, and CNN2 transcripts (**Figure 2C**).

**Figure 2.**
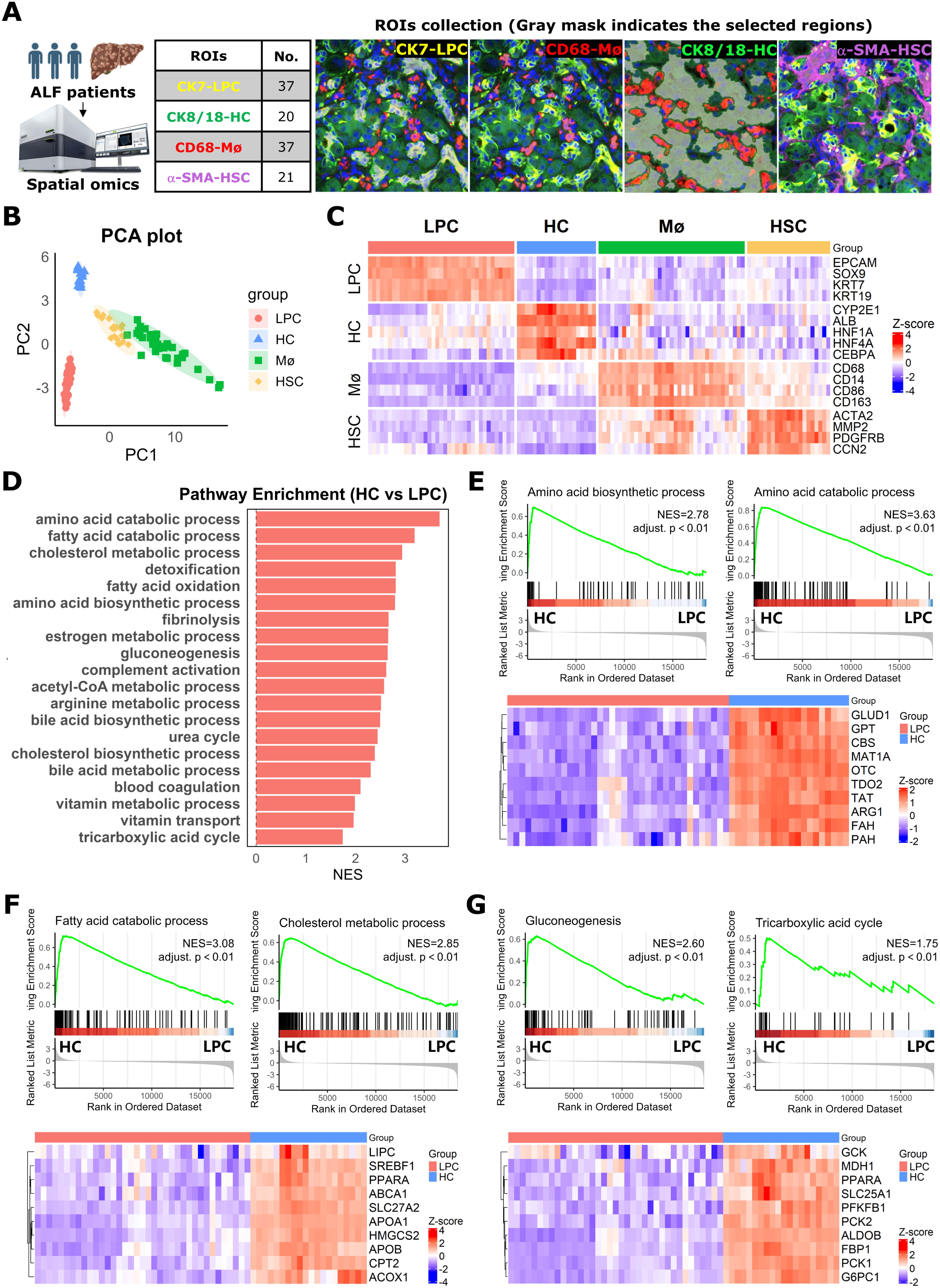
Spatial transcriptomic analysis reveals the transcriptomes of liver progenitor cells, hepatocytes, macrophages, and hepatic stellate cells. **(A)** A scheme depicts the GeoMx DSP workflow. A total of 115 regions of intrest (ROIs) were collected based on cell-type-specific markers: CK7⁺ LPCs (yellow, n = 37), CK8/18⁺ HCs (greeen, n = 20), CD68⁺ Macrophages (Mø) (red, n = 37), and α-SMA⁺ HSCs (violet, n = 21). Gray-masked regions indicate the precise ROIs selected for transcriptomic profiling within each cell type. **(B)** Principal component analysis (PCA) based on whole-transcriptome profiles shows distinct clustering of LPCs, HCs, Macrophages, and HSCs. **(C)** Heatmap illustrates representative cell-type-specific gene expression patterns. **(D)** GO BP pathway enrichment analysis was performed to identify enrichments of metabolic function genes between HCs and LPCs, including pathways for amino acid, fatty acid, and cholesterol metabolism, detoxification, and gluconeogenesis. **(E–G)** GSEA and corresponding heatmaps reveal transcriptional downregulation of hepatic functional pathways in LPCs, including amino acid metabolism, lipid metabolism, gluconeogenesis and the tricarboxylic acid cycle. Representative metabolic genes (e.g., GLUD1, PPARA, FBP1, PCK1, et al) are listed.

Gene Ontology Biological Process (GO BP) analysis revealed distinct functional pathways between hepatocytes and LPCs, including amino acid and fatty acid catabolic processes, cholesterol, estrogen, acetyl-CoA, arginine, bile acid and vitamin metabolic processes; amino acid, bile acid and cholesterol biosynthesis processes; detoxification, fibrinolysis, gluconeogenesis; complement activation; the urea cycle; blood coagulation; vitamin transport and tricarboxylic acid (TCA) cycle (**Figure 2D**). Gene set enrichment analysis (GSEA) showed significantly reduced expression of genes relevant to above functional pathways in LPCs compared to hepatocytes (**Figure 2E-G**). As shown in **Figure 2E-G**, genes associated with the biosynthesis and catabolism of amino acids (e.g., GLUD1, GPT, CBS, and TAT), fatty acids (e.g., PPARA, ACOX1, and CPT2), and cholesterol (e.g., SREBF1, ABCA1, and HMGCS2), as well as gluconeogenesis (e.g., G6PC1, PCK1, and FBP1) and TCA cycle (e.g., MDH1 and SLC25A1), were markedly reduced in LPCs. Additional significantly reduced genes in LPCs were associated with coagulation factors (e.g., F2, F7, F9, and F10); detoxification factors (e.g., GSTA1, PRDX1, and GSTO1); the estrogen metabolic process (e.g., UGT2B7, HSD17B1, and CYP2C9); the bile acid metabolic process (e.g., CYP27A1, ABCB11, and SULT2A1); the urea cycle (e.g., CPS1, ARG1, and OTC), and vitamin transport (e.g., SLC23A2, SLC22A1, and TTPA) (**Figure S1**).

These pathways are hallmarks of mature liver function and highlight the transcriptional immaturity of LPCs. These results demonstrate the transcriptome changes that occur during the transition from LPCs to mature hepatocytes.

### Cellular communications between liver progenitor cells and surrounding cells macrophages, hepatic stellate cells and hepatocytes in acute liver failure

Subsequently, we analyzed the cellular communications between LPCs, remaining hepatocytes, macrophages and activated HSCs in these irreversible ALF patients. As shown in **Figure 3A**, there were high levels of cell-cell interaction counts and weights between LPCs, macrophages and HSCs. On the one hand, LPCs received multiple strong signals from surrounding cells, including TGF-β, activin, EGF, HGF, FGF, TWEAK and IGF (**Figure 3B**). On the other hand, LPCs also sent strong signals to surrounding cells, including TGF-β, EGF, FGF, CXCL, CSF, and PDGF (**Figure 3B**). Further cell-cell communication analysis revealed signal transduction direction between LPCs and surrounding cells. Macrophages and activated HSCs sent TGF-β, BMP, NODAL, EGF, HGF, FGF, TWEAK, IGF and CXCL to LPCs (**Figure 3C**). LPCs sent TGF-β, activin, BMP, NODAL, EGF, HGF, FGF, CSF, PDGF and CXCL to macrophages and TGF-β, activin, BMP, NODAL, EGF, FGF, CSF, PDGF and CXCL to activated HSCs (**Figure 3C**). In addition, LPCs produced TGF-β, activin, BMP, EGF, FGF, TWEAK, IGF and CXCL for autoregulation (**Figure 3C**). Further ligand-receptor analyses showed that the macrophage- and HSC-derived TGF-β1-ACVR1-TGFBR1 signaling pathway was a predominant signal received by LPCs (**Figure 3D-E**), suggesting a potential critical role of this signaling pathway in regulating LPC behaviors in ALF.

**Figure 3.**
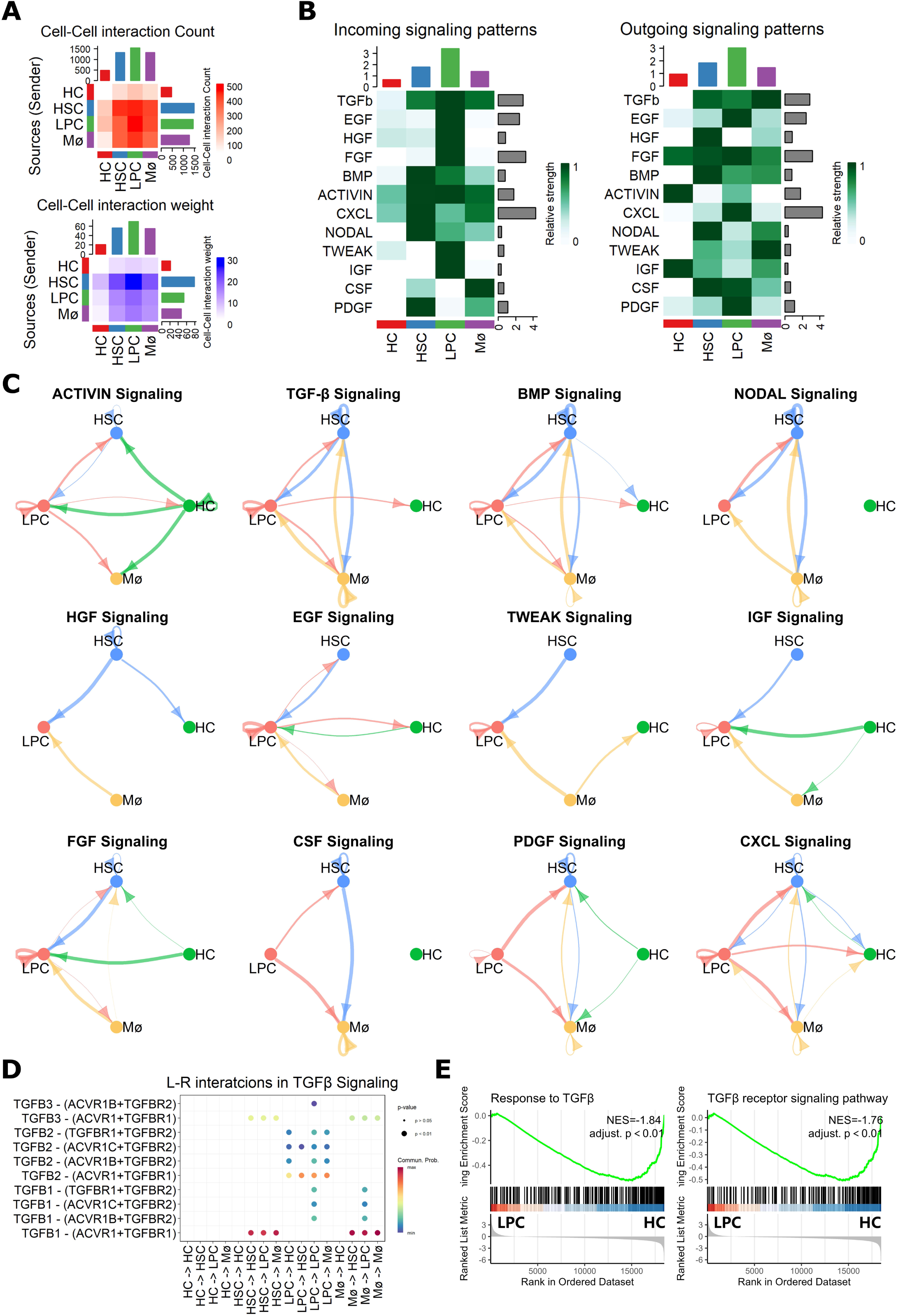
Cell communication between liver progenitor cells, macrophages and hepatic stellate cells. **(A)** Cell–cell interaction analysis based on spatial transcriptomic data was performed to evaluate interaction counts and interaction weights between LPCs, macrophages (Mø), and HSCs. **(B)** Heatmaps summarize the incoming (left) and outgoing (right) signaling patterns in LPCs, Mø and HSCs. **(C)** Network diagrams illustrate the directionality and intensity of specific signaling pathways among LPCs, HSCs, Mø, and HCs. Line thickness indicates interaction strength, and arrow direction denotes the source-to-target relationship. **(D)** Dot plots display specific ligand-receptor (L-R) interactions in TGFβ signaling pathways between different cell populations. **(E)** Gene set enrichment analysis (GSEA) highlights enrichment of “Response to TGF-β” and “TGF-β receptor signaling pathway” gene sets in LPCs compared to HCs.

### TGF-β controls LPC proliferation

The potential importance of the TGF-β signaling pathway has also been demonstrated in a recent spatiotemporal analysis of mice fed with 3,5-diethoxycarbonyl-1,4-dihydrocollidine (DDC)^26^. In DDC-fed mice, TGF-β signaling is a predominant pathway regulating DDC-induced DRs and liver tissue recovery after the cessation of DDC toxicity^26^. Given the key role of TGF-β in cell proliferation^15^, we examined liver tissues collected from wild-type and SMAD7 transgenic mice fed with DDC for 4 weeks. SMAD7 is a negative feedback regulator, to end TGF-β action^27^.

Compared to DDC-fed wild-type mice, p-SMAD3 expression in SMAD7 transgenic mice was remarkably inhibited (**Figure 4A)**, suggesting the silence of TGF-β signaling in these mice. Concomitant with the inhibition of TGF-β signaling, IHC and IF staining showed that CK19- and bromodeoxyuridine (BrdU)-marked LPC proliferation was significantly increased in SMAD7 transgenic mice compared to wild-type controls (**Figure 4B**). To quantify these data, we performed integrated optical density (IOD) analysis of CK19 staining per unit area in all mice (**Figure S2A**). The quantification data demonstrated that SMAD7 transgenic mice exhibited approximately a twofold increase in CK19 expression compared to wild-type controls (11.624 ± 1.88 vs 6.228 ± 1.23, *p* < 0.0001) (**Figure S2B**). These results suggest that TGF-β acts as an anti-proliferative regulator in LPC proliferation.

**Figure 4.**
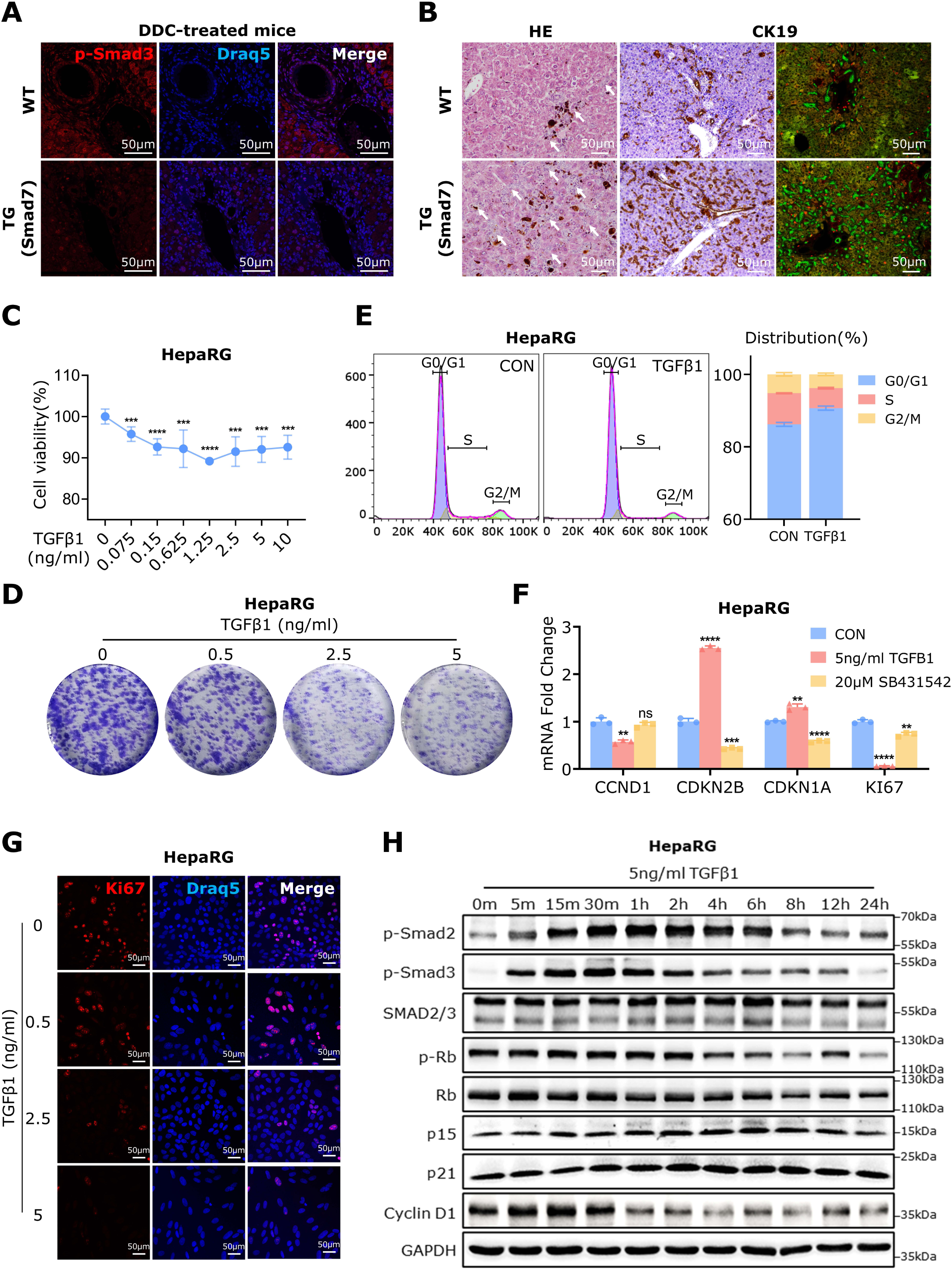
TGF-β controls LPC proliferation. **(A)** Immunofluorescence (IF) staining for p-Smad3 was performed on liver tissues from wild-type (WT) and SMAD7 transgenic (TG) mice fed with DDC for 4 weeks. **(B)** Hematoxylin and eosin (H&E) staining, immunohistochemistry (IHC) and IF for CK19 (green) and bromodeoxyuridine (BrdU, red) were performed on liver tissues collected from mice fed with DDC for 4 weeks. Mice received BrdU injection 2 h before sacrifice. Arrows in the H&E staining highlight ductular reaction areas. **(C)** Cell viability was analyzed in HepaRG cells treated with different concentrations of TGF-β1 for 48h using the CCK-8 assay. **(D)** A colony formation assay was performed on HepaRG cells exposed to various concentrations of TGF-β1 for 14 days. **(E)** Flow cytometric analysis was conducted to examine the cell cycle distribution in HepaRG cells treated with or without TGF-β1 for 24h. **(F)** Quantitative reverse transcription PCR (qRT-PCR) was used to analyze mRNA expression of CCND1, CDKN2B, CDKN1A, and KI67 in HepaRG cells treated with TGF-β1 and SB431542. **(G)** IF staining for Ki67 was performed on HepaRG cells treated with TGF-β1 at indicated concentrations. **(H)** Western blotting was conducted to analyze SMAD and cell cycle-related proteins in HepaRG cells treated with 5 ng/mL TGF-β1 at different time points. Scale bars: 50 μm (A, B, G). Data are presented as mean ± SD. *p < 0.01 (**), p < 0.001 (***), p < 0.0001 (****)*; ns, not significant.

To further investigate how TGF-β1 controls LPC proliferation, we performed a CCK-8 assay in the human LPC line HepaRG cells, which were treated with different concentrations of TGF-β1. As shown in **Figure 4C**, TGF-β1 dose-dependently reduced LPC cell viability. A colony formation assay also demonstrated that TGF-β1 dose-dependently decreased LPC colony formation (**Figure 4D**). Flow cytometric (FACS) analysis further revealed that TGF-β1 arrested LPC cells in the G1 phase, preventing their entry into the S phase (**Figure 4E**). Consistent with the FACS finding, qRT-PCR analysis showed that TGFβ1 inhibited the mRNA expression of CCND1 while increasing the expression of CDKN2B and CDKN1A (**Figure 4F**). Furthermore, both mRNA and protein expression of KI67 were significantly inhibited in TGF-β1-treated HepaRG cells (**Figure 4F-G**). The effects of TGF-β1 were reversed by the TGF-β receptor I inhibitor SB431542 (**Figure 4F**).

Next, we performed Western blot analysis to investigate the dynamic alterations of cell cycle regulatory proteins in HepaRG cells under TGF-β1 stimulation. TGF-β1-induced p-SMAD2 and p-SMAD3 levels rapidly increased within 5 minutes of stimulation and remained high until 24 hours (**Figure 4H**). TGF-β1 stimulation reduced the expression of Cyclin D1 and p-Rb over time while rapidly increasing the expression of p15 and p21, indicating TGF-β1-induced cell cycle arrest (**Figure 4H**). These results suggest that TGF-β1 plays a decisive role in determining the G1 to S phase transition in LPCs, similar to its role in other cells^28^.

### Active TGF-β signaling in proliferating liver progenitor cells

To clarify the activity of TGF-β signaling in LPCs, we performed IHC to examine the expression of p-SMAD2 in 8 irreversible ALF patients. As shown in **Figure 5A**, robust p-SMAD2 expression was observed in LPCs, macrophages, and HSCs, indicating active TGF-β signaling in these cells. Intriguingly, these cells, including LPCs, simultaneously expressed high levels of c-MYC (**Figure 5A, Figure S3A**), suggesting that TGF-β signaling does not inhibit LPC proliferation in ALF following MHN.

**Figure 5.**
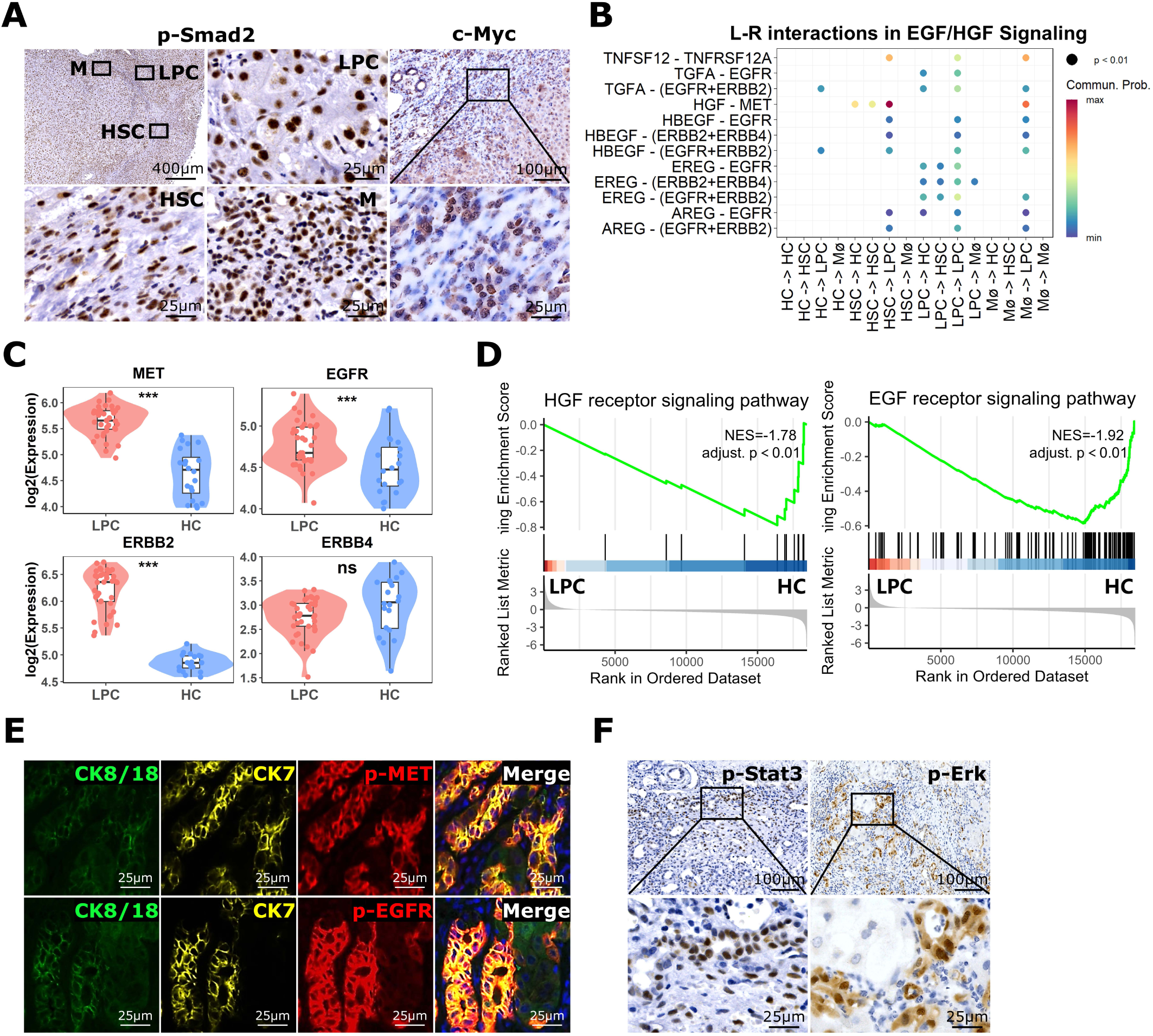
TGF-β, HGF, and EGF signaling are active in proliferating LPCs. **(A)** Immunohistochemistry (IHC) staining for p-SMAD2 and c-MYC was performed in liver tissues from ALF patients. Frames highlight p-SMAD2 expression in LPCs, HSCs, and macrophages (Mø). **(B)** Ligand–receptor interaction analysis focusing on EGF and HGF signaling pathways was conducted to predict communication intensity between LPCs, HSCs, Mø, and hepatocytes (HCs). **(C)** Violin plots display expression levels of MET, EGFR, ERBB2, and ERBB4 in LPCs and HCs. **(D)** GSEA was performed to compare HGF and EGF receptor signaling pathways between LPCs and HCs. **(E)** Immunofluorescence (IF) staining for CK8/18 (green), CK7 (yellow), and phosphorylated EGFR (p-EGFR) or phosphorylated MET (p-MET) (red) was conducted on liver tissues from ALF patients. Scale bar: 25 μm. **(F)** IHC staining of p-STAT3 and p-ERK was conducted on liver tissues from ALF patients. Scale bar: 25 and 100 μm.

How can LPCs proliferate in the presence of active TGF-β-SMAD signaling? Spatial transcriptome and ligand-receptor analysis showed that, in addition to TGF-β, LPCs received multiple proliferative signals from surrounding cells, including TNFSF12 (TWEAK), TGF-α, HGF, HBEGF, EREG, and AREG (**Figure 3B**). Among these signals, macrophage- and HSC-derived HGF-MET signaling provided the strongest proliferative stimulus for LPCs (**Figure 5B**). GSEA analysis revealed a remarkable increase in the expression of HGF and EGF receptor genes (e.g., MET, EGFR, ERBB2and ERBB4) in LPCs compared to hepatocytes (**Figure 5C-D**). The substantial expression of p-MET and p-EGFR in LPCs, but not in hepatocytes, was confirmed in the same four ALF patients through Co-IF staining. Subsequently, we performed IHC for p-STAT3 and p-ERK in these patients. As shown in **Figure 5E** and **Figure S3B**, both p-STAT3 and p-ERK were positively expressed in LPCs. These findings suggest that both HGF and EGF signaling pathways are activated in LPCs.

### HGF promotes proliferation of liver progenitor cells with TGF-β stimulation

Compared to healthy individuals, serum concentrations of HGF in patients with fulminant hepatic failure were remarkably increased (from 0.27 ± 0.08ng/mL to 16.40 ± 14.67ng/mL)^29^. Therefore, we examined the effects of HGF on LPC proliferation.

CCK-8 assay showed that HGF dose-dependently increased the cell viability of HepaRG cells (**Figure 6A**). To investigate how HGF drives LPC proliferation, we stimulated LPCs with HGF for different durations (e.g., 1 h, 8 h, and 24 h) and performed a time-resolved transcriptomic analysis (**Figure 6B**). Unsupervised clustering revealed six distinct gene expression modules (C1, C2, C3, C4, C5, and C6) after dynamic HGF stimulation (**Figure 6B**): (1) Module C1 comprises 4207 genes, which reached peak expression at 8 h after HGF stimulation; (2) Module C2 contains 4209 genes, which were downregulated following HGF stimulation for 8 h and 24 h; (3) Module C3 consists 3842 genes, which were inhibited by HGF stimulation for 1h and 8 h and then restored to baseline levels at 24 h post stimulation; (4) Module C4 comprises 3558 genes, which remains stable after 1 h and 8 h of HGF stimulation but were upregulated after 24 h of treatment; (5) Module C5 includes 2863 genes, which were upregulated by HGF stimulation for 1 h and subsequently downregulated to normal levels; and (6) Module C6 has 2860 genes, which exhibits stable expression after 1 h and 8 h of HGF stimulation but were downregulated at 24 h. Gene set variation analysis (GSVA) further showed that transient activation of the MYC target gene signature was induced by HGF stimulation for 1 h and peaked at 8 h. In contrast, the E2F target gene signature reached its peak 8 h after HGF stimulation and remained elevated until 24 h (**Figure 6C**).

**Figure 6.**
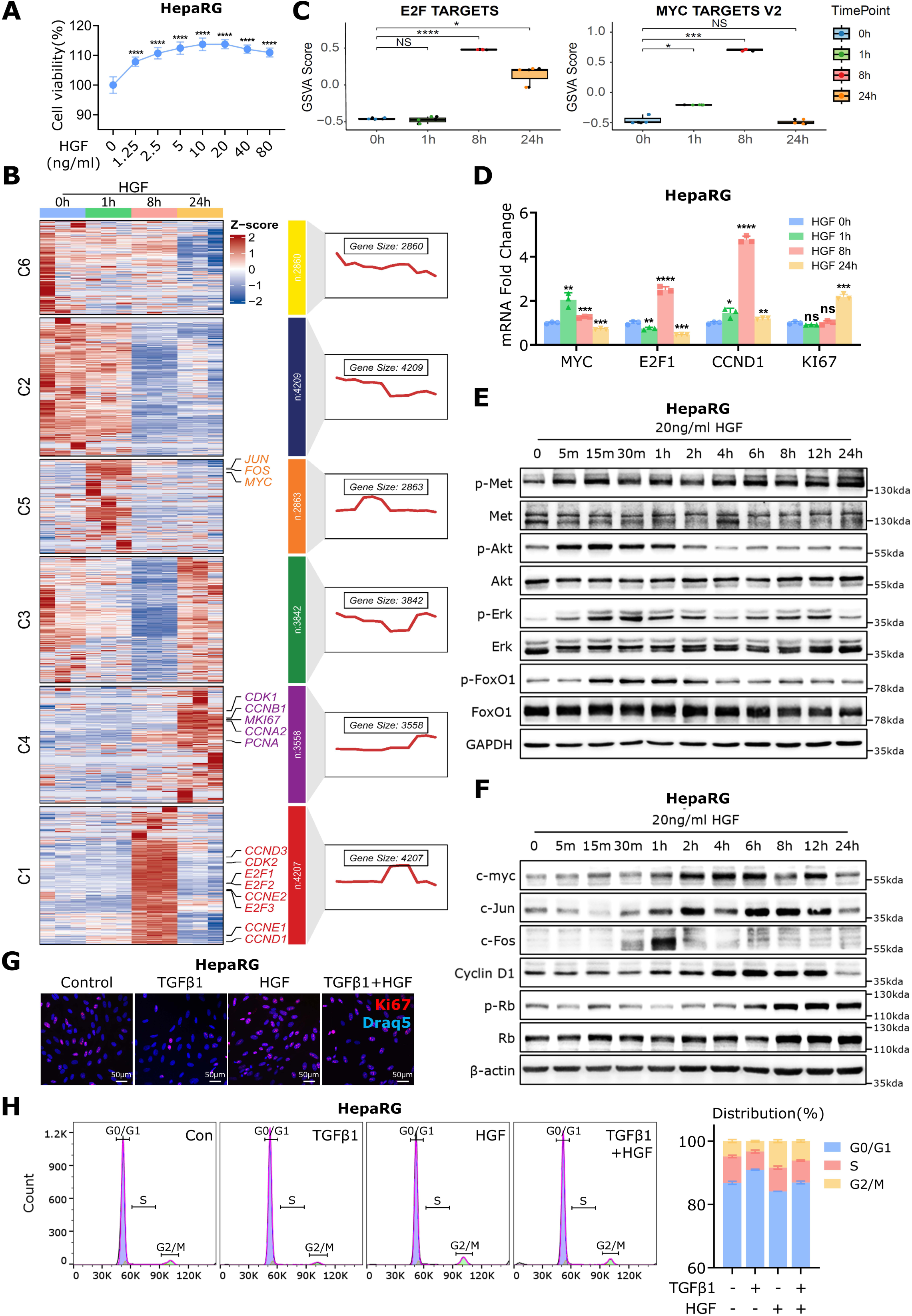
HGF overrides TGF-β and promotes LPC proliferation. **(A)** Cell viability was assessed in HepaRG cells treated with different concentrations of HGF for 48h using the CCK-8 assay. **(B)** A heatmap displays transcriptomic changes in HepaRG cells treated with 20 ng/mL HGF for 0, 1, 8, and 24 h. Genes were grouped into six clusters (C1–C6) based on distinct expression patterns. Altered cell cycle– and transcription factor–related genes are annotated within each cluster. Corresponding enrichment curves and gene counts illustrate dynamic expression trends over time. **(C)** Gene Set Enrichment Analysis (GSVA) scores were calculated to assess E2F and MYC target gene sets at indicated time points. **(D)** qRT-PCR was performed to examine mRNA expression of MYC, E2F1, CCND1, and KI67 in HepaRG cells treated with HGF for the indicated durations. **(E)** Western blot analysis was conducted to assess phosphorylated and total Met, Akt, Erk, and FoxO1 in HepaRG cells treated with 20 ng/mL HGF for the indicated time points. **(F)** Western blotting was used to analyze protein expression of c-Myc, c-Jun, c-Fos, CCND1, phosphorylated and total Rb in HepaRG cells stimulated with 20 ng/mL HGF for different time periods. **(G)** Immunofluorescence (IF) staining for Ki67 and Draq5 was performed in HepaRG cells treated with TGF-β1, HGF or a combination of both. Scale bar: 50 μm. **(H)** Flow cytometry was used to analyze cell cycle distribution in HepaRG cells under TGF-β1, HGF and TGF-β1+HGF treatment. The right panel summarizes the percentage of cells in the G0/G1, S, and G2/M phase. Data are presented as mean ± SD. *p < 0.05 (*), p < 0.01 (**), p < 0.001 (***), p < 0.0001 (****)*; ns, not significant.

Subsequent qPCR assay revealed that HGF stimulation for 1 h significantly increased mRNA expression of c-MYC (**Figure 6D**). E2F1 and CCND1 expression was upregulated after 8 h of HGF stimulation (**Figure 6D**). HGF incubation for 24 h resulted in high levels of Ki67 mRNA (**Figure 6D**). Subsequently, we performed Western blotting to examine the dynamical expression of HGF downstream signaling molecules and targeted proteins in HepaRG cells. HGF administration for 5 minutes was sufficient to induce high levels of expression of p-MET, p-AKT and p-ERK in LPCs (**Figure 6E**). p-FOXO1 expression was induced after 15 minutes of HGF incubation (**Figure 6E**). The peak levels of p-MET persisted until 24 h, whereas p-AKT and p-FOXO1 expression was maintained for only 2 h (**Figure 6E**). p-ERK expression lasted up to 12 h (**Figure 6E**). Administration of HGF significantly induced the expression of c-MYC, c-Jun, CCND1, and p-RB in LPCs (**Figure 6F**). Expression of c-MYC, c-Jun and CCND1 were robustly upregulated after 1 h of HGF incubation and lasted for 12 h (**Figure 6F**). p-RB expression was upregulated after 6 h of HGF incubation and persisted until 24 h (**Figure 6F**).

IF staining further revealed that HGF increased Ki-67 expression in HepaRG cells even in the presence of TGFβ1 stimulation (**Figure 6G**). FACS assay further demonstrated that HGF promoted G1-to-S phase progression in HepaRG cells, regardless of TGF-β stimulation (**Figure 6H**). These results suggest that HGF largely compromises the ability of TGFβ1 to arrest LPCs from entering the S phase from the G1 phase.

### EGF promotes liver progenitor cell proliferation at the presence of TGF-β

In addition to HGF, we investigated how EGF regulates LPC proliferation. CCK-8 and colony formation assays showed that EGF dose-dependently increased cell viability and colony formation in HepaRG cells (**Figure 7A-B**). qRT-PCR analysis further revealed that 1-hour EGF stimulation sufficiently induced high levels of c-MYC mRNA expression (**Figure 7C**). EGF incubation also continuously increased CCND1 mRNA expression in LPCs (**Figure 7C**). Dynamical Western blot analysis showed that EGF administration for 5 minutes rapidly induced the expression of p-ERK, p-AKT, and p-FOXO1 (**Figure 7D**). The peak levels of these proteins persisted for up to 1 hour (**Figure 7D**). EGF downstream transcription factor c-Fos was induced 30 minutes after EGF stimulation and remained elevated for up to 2 hours (**Figure 7D**). Additionally, EGF stimulation for 5 minutes was sufficient to induce high levels of the expression of c-MYC, p-RB, and Cyclin D1, with these effects lasting for more than 12 hours (**Figure 7D**). ChIP-qPCR analysis further demonstrated that c-Myc binds to the CCND1 promoter (-247bp ∼ -31bp and -797bp ∼ -593bp) upon EGF stimulation in HepaRG cells (**Figure 7E**).

**Figure 7.**
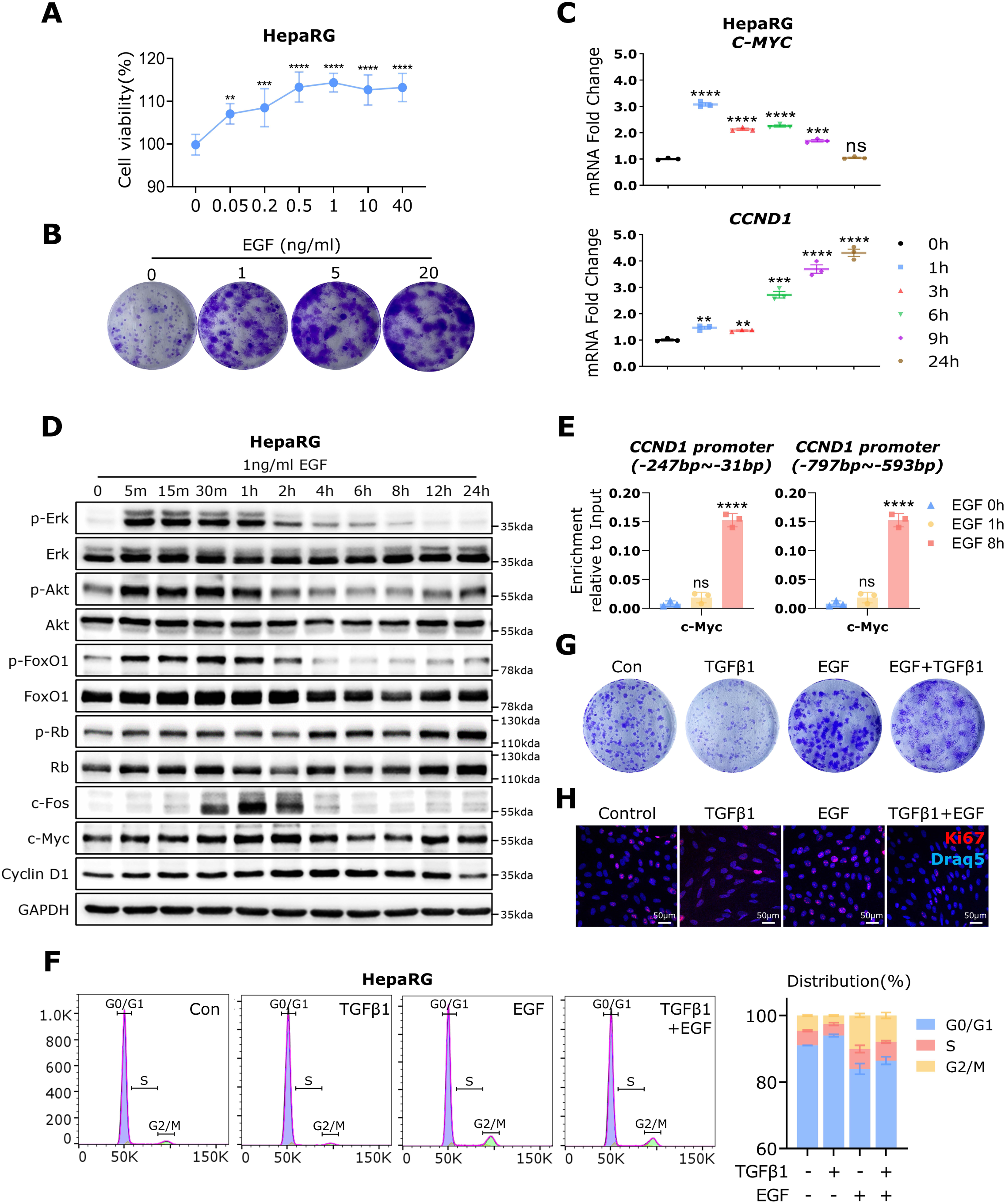
EGF overrides TGF-β and promotes LPC proliferation. **(A)** Cell viability was assessed in HepaRG cells treated with different concentrations of EGF for 48 h using the CCK-8 assay. **(B)** A colony formation assay was performed in HepaRG cells treated with the indicated concentrations of EGF for 14 days. **(C)** qPCR was used to measure mRNA expression of C-MYC and CCND1 in HepaRG cells treated with 1 ng/mL EGF at 0, 1, 3, 6, 9 and 24 h. **(D)** Western blotting was perfomed to examine phosphorylated and total ERK, AKT, FoxO1, Rb, c-Fos, c-Myc, and Cyclin D1 in HepaRG cells treated with 1 ng/mL EGF at the indicated time points. **(E)** ChIP-qPCR was used to analyse c-Myc binding to two promoter regions of the CCND1 gene (∼247 bp to ∼31 bp and ∼797 bp to ∼593 bp) in HepaRG cells with EGF treatment for 8 h. **(F)** Flow cytometry was performed to analyze cell cycle distribution in HepaRG cells treated with TGF-β1 (5 ng/mL), EGF (1 ng/mL), or both for 24 h. The right panel summarizes the percentage of cells in the G0/G1, S, and G2/M phase. **(G)** A colony formation assay was performed in HepaRG cells treated with TGF-β1, EGF and TGF-β1+EGF for 14 days. **(H)** Immunofluorescence (IF) staining for Ki67 and Draq5 was performed in HepaRG cells under the TGF-β1, EGF and TGF-β1+EGF 24h treatment. Scale bars: 50 μm. Data are presented as mean ± SD. *p < 0.01 (**), p < 0.001 (***), p < 0.0001 (****)*; ns, not significant.

FACS assay further demonstrated that EGF promoted G1-to-S phase progression in HepaRG cells (**Figure 7F**). In the presence of EGF, the ability of TGFβ1 to arrest LPCs from entering the S phase was largely compromised (**Figure 7F**). Consistent with the FACS analysis, a colony formation assay showed that EGF incubation significantly promoted cell proliferation, even in the presence of TGFβ1 (**Figure 7G**). Furthermore, IF staining revealed that EGF increased Ki-67 expression in HepaRG cells treated with TGFβ1 (**Figure 7H**). These results suggest that EGF counteracts the anti-proliferative effect of TGFβ on LPCs.

### Activated SMADs are essential for HGF-dependent HNF4α transcription in LPCs

It has been shown that HGF and EGF exert opposing effects in regulating LPC phenotype in mice fed with DDC^30^. EGF promotes cholangiocyte specification and suppresses LPC differentiation into hepatocytes. In contrast, HGF induces LPCs to express hepatocyte phenotypes^30^.

To examine the effects of HGF on the expression of hepatocyte and cholangiocyte genes in LPCs, we performed qPCR analysis in HepaRG cells stimulated with HGF for different duration. In contrast to the previous study^30^, HGF inhibited the mRNA expression of hepatocyte genes, such as HNF1α, HNF4α, and albumin, while increasing the expression of SOX9 and keratin 19 (**Figure S4**). Notably, HepaRG cells are a type of hepatic bipotent progenitor cell line derived from normal hepatocytes surrounding tumor tissues from a patient with cholangiocarcinoma and HCV infection^31,32^. These LPCs maintain multiple liver function genes, such as CYP enzymes and transporters^32^. In contrast to hepatocyte-originated LPCs, such as HepaRG cells, activated LPCs in MHN-associated ALF are derived from smallest cholangiocytes located in the Canal of Hering^1^. To confirm the effects of HGF on cholangiocyte-derived LPCs, we stimulate biopotential murine oval liver (BMOL) cells with HGF^33^.

Consistent with the previous study^30^, qPCR analysis in this study showed that HGF induced the mRNA expression of HNF4α and HNF1α while inhibiting the expression of SOX9 in the murine LPC line BMOL cells (**Figure 8A**). Interestingly, HGF-dependent upregulation of HNF4α and HNF1α expression was inhibited when LPCs were treated with the general TGFβ receptor inhibitor SB431542 (**Figure 8B**). SB431542 also reduced SOX9 mRNA expression in BMOL cells (**Figure 8B**). Subsequently, we used RNAi to knock down the expression of either ALK5 or SMAD3. Knockdown of SMAD3 remarkably reduced the mRNA expression of CPS1, albumin, C/EBPα, HNF1α, and HNF4α, while knockdown of ALK5 inhibited the mRNA expression of CPS1, albumin and C/EBPα (**Figure 8C**). Disruption of either SMAD3 or ALK5 did not alter SOX9 expression (**Figure 8C**).

**Figure 8.**
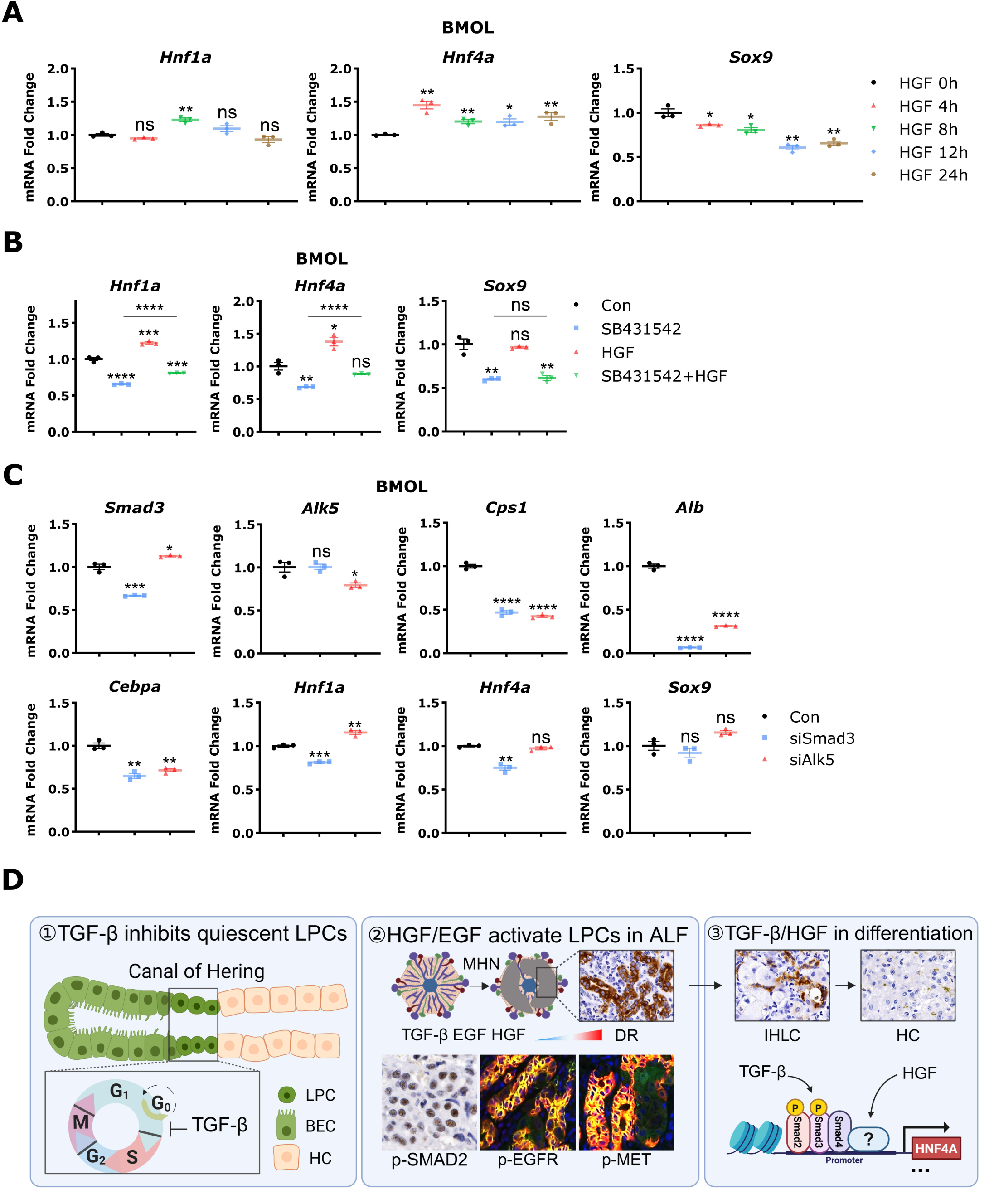
TGF-β signaling is essential for HGF-induced HNF4α expression. **(A)** qRT-PCR was used to measure mRNA expression of Hnf1a, Hnf4a, and Sox9 in BMOL cells treated with 20 ng/mL HGF for 0, 4, 8, 12 and 24 h. **(B)** qRT-PCR was performed to analyze mRNA expression of Hnf1a, Hnf4a, and Sox9 in BMOL cells treated with SB431542 (20 μM) for 48 h, followed by HGF (20 ng/mL) for an additional 8 h. **(C)** qRT-PCR was used to analyze mRNA expression of Smad3, Alk5, Hnf1a, Hnf4a, Cebpa, Cps1, albumin, and Sox9 in BMOLs incubated with siRNA targeting Smad3 or Alk5 incubation for 48 h. **(D)** A graphic abstract illustrates: (1) TGF-β–mediated inhibition of quiescent LPCs, (2) Activation of LPCs by HGF/EGF signaling during massive hepatic necrosis (MHN), and (3) Coordinated roles of TGF-β and HGF in LPC differentiation toward hepatocytes by regulating transcriptional factors such as HNF4A. Data are shown as mean ± SD. *p < 0.05 (*), p < 0.01 (**), p < 0.001 (***), p < 0.0001 (****)*; ns, not significant.

## Discussion

In the current study, we performed a spatial transcriptome analysis to investigate the critical microenvironmental signals that control LPC proliferation and performing hepatocyte functions in MHN-associated ALF. Among multiple essential signal mediators, TGF-β plays a unique role in regulating LPC behavior. We found the following: (1) As in hepatocytes^18^, TGF-β inhibits LPC proliferation both *in vitro* and in DDC-fed mice. Its anti-proliferative effect is achieved through SMAD-mediated expression of the CDK inhibitors p15 and p21, which suppress cyclin D1 expression and arrest cell cycle transition from the G1 to the S phase. (2) In ALF patients with MHN, robustly proliferating LPCs exhibit strong p-SMAD2/3 expression, suggesting that additional powerful signals override the anti-proliferative effect of TGF-β and drive LPC proliferation. (3) According to spatial transcriptome analysis, proliferating LPCs receive abundant signals from surrounding macrophages and HSCs, including complete mitogens HGF and EGF. Both HGF and EGF are capable of inducing LPC proliferation in the presence of TGF-β. (4) In addition to inducing LPC proliferation, HGF promotes the expression of multiple vital hepatocyte genes (e.g., HNF4α and HNF1α) in LPCs. These results illustrate the central role of TGF-β in regulating LPC behaviors.

It is not known yet how LPCs are kept in a quiescent state physiologically. This study revealed that TGF-β signaling might serve as a key regulator preventing LPC activation in normal conditions. We found that TGF-β impedes LPC cell cycle progression from the G1 to the S phase by upregulating the expression of the genes CDKN1A and CDKN2B, which encode the two CDK inhibitors p15^INK4B^ and p21^Cip1^, respectively. Additionally, TGF-β also represses c-MYC expression in LPCs. As in other cell types^34^, the anti-proliferative effect of TGF-β in LPCs is reversible. The most robust LPC proliferation occurs in MHN-associated ALF, which is defined as a type III ductular reaction by Desmet^7^. In this study, spatial transcriptome analysis revealed that LPCs receive multiple critical signals from surrounding macrophages and HSCs, including anti-proliferative mediators such as TGF-β and pro-proliferative factors like HGF and EGF. The activation of HGF-p-MET-p-STAT3 and EGF-p-EGFR-p-ERK signaling pathways in LPCs of ALF patients was confirmed by IHC and IF staining. Notably, robust expression of p-MET, p-STAT3, p-EGFR, and p-ERK was mainly observed in LPCs, while absent in the remaining hepatocytes in these ALF patients, suggesting that LPCs are primary cells mediating liver regeneration in this scenario. In vitro studies further demonstrated that both HGF and EGF are sufficient to induce LPC proliferation, even in the presence of high concentrations of TGF-β. These findings explain why p-SMAD2 positive LPCs still undergo robust proliferation: HGF and EGF override the inhibitory signals to drive cell cycle progression in this setting. In a cohort of ALF patients, HGF concentrations reached 16.4 ± 14.67 ng/mL—more than 60 times the levels observed in healthy controls (0.27 ± 0.08 ng/mL)^29^. Therefore, it is not surprising that LPCs predominantly received HGF stimulation rather than EGF in MHN-associated ALF. According to our spatial transcriptome analysis and previous studies^35,36^, macrophages are one of the main sources of HGF in ALF.

An interesting question is what role the TGF-β-SMAD signaling pathway plays when its anti-proliferative effect on LPCs is compromised in MHN-associated ALF. We observed that, in addition to functioning as a complete mitogen, HGF is a critical growth factor inducing hepatocyte genes such as HNF4α in LPCs^30^. According to our previous study, transcription of HNF4α in LPCs requires p-SMADs^37^. In hepatocytes, constitutive HNF4α expression also depends on p-SMAD proteins to recruit the acetyltransferase CREB-binding protein/p300^21^. Although we did not directly examine this mechanism in LPCs, it is conceivable that a similar SMAD–CBP/p300 interaction facilitates HGF-induced HNF4α transcription in LPCs. These findings highlight an essential role of the TGF-β-SMAD signaling pathway in MHN-associated ALF.

In a normal liver, the maintenance of cholangiocyte phenotype is associated with TGF-β2 and TGF-β3^19^. However, in the context of MHN-associated ALF, spatial transcriptomic analysis shows that LPCs primarily receive TGF-β1 signal, rather than TGF-β2 or TGF-β3. Although HSCs are traditionally considered the major source of TGF-β1 in liver diseases^38,39^, cell–cell communication analysis revealed that macrophages, rather than fibroblasts, are the predominant source of TGF-β1 in ALF patients (**Figure 3C**). Activated macrophages are the major source of TGF-β1 in this setting. This shift in the source of TGF-β1 from HSCs to macrophages may reflect a unique inflammatory microenvironment in MHN-associated ALF. How macrophage-produced TGF-β1 affects inflammatory microenvironment in this disease scenario is worth further investigation.

Taken together, this study highlights a critical role of TGF-β in regulating LPCs both in normal and pathological situations. As summarized in **Figure 8D**, TGF-β maintains LPCs in a quiescent state in a normal liver. Once ALF occurs, MHN induces severe inflammatory cell infiltration (e.g., macrophage recruitment) to clear dead hepatocytes and restoring local homeostasis, HSC activation for wound repair, and LPC proliferation to restore liver function. Proliferating LPCs produce chemokines that recruit macrophages and activated HSCs. Macrophages produce substantial amount of HGF, which further stimulates LPC proliferation and differentiation toward hepatocytes in collaboration with TGF-β. In this scenario, TGF-β serves as an integrative signal that determines LPC functions and fate.

## Materials and Methods

### Patients

A total of 8 ALF patients were enrolled in the Department of Gastroenterology and Hepatology, Beijing You’an Hospital, Affiliated with Capital Medical University between 2015 and 2018. All patients were diagnosed with HBV-induced ALF and the liver tissue were collected when patients received liver transplantation. These patients have been investigated in previous studies^37,40^. Tissue samples were fixed in 10% formalin for 24 hours and embedded in paraffin for histological measurement, immunofluorescence, immunohistochemistry, and GeoMx digital spatial profiler analysis.

In this study, ALF is defined as severe acute liver injury lasting more than 4 weeks duration with coagulation abnormalities (INR≥1.5) and any degree of mental alteration (encephalopathy) in a patient without preexisting liver disease. The study protocol was approved by local ethics committees (Jing-2015-084). Written informed consent was obtained from the patients or representatives.

### Animals

SMAD7 transgenic mice were described previously^41^. SMAD7 transgenic mice and their wild-type littermates (8-10 weeks old) were fed by a diet containing 3,5-diethoxycarbonyl-1,4-dihydrocollidine (DDC, 0.1% wt/wt in Purina 5015 mouse chow; Bio-Serv, Frenchtown, NJ) for 4 weeks to induce LPC proliferation^42^. Mice received BrdU injection 2 h before sacrifice. Liver samples from multiple lobes were fixed in 4% buffered formalin for histological analysis. The animal experiments were approved by local institutional animal care committee and conformed to the local guidelines for animal care.

### Geomx digital spatial profiler

Spatial transcriptomics was performed by GeoMx digital spatial profiler (NanoString) with the probes of whole transcriptome atlas panel. Formalin-Fixed Paraffin-Embedded (FFPE) tissue sections (5 μm) were baked at 60°C for 30 minutes. Subsequently, slides were deparaffinized by xylene, underwent antigen retrieval in Tris-EDTA buffer (pH 9.0, 00-4956-58; Thermo Fisher Scientific) for 20 minutes and incubated by 1 μg/mL proteinase K (AM2546; Thermo Fisher Scientific) treatment for 15 minutes at 37°C, respectively. After fixation with 10% neutral buffered formalin (15740-04; EMS Diasum) for 5 minutes and RNA probe hybridization at 37°C overnight, sections were stringently washed and stained with morphology markers (CK8/18-AF488, CK7-AF594, CD68-AF647) and nuclear stain SYTO83-CY3 for 1 hour. Based on the fluorescence signals, we defined and selected four phenotypically distinct regions of interests (ROI): CK7^+^ LPCs (n=37), CK8/18^+^ HCs (n=20), α-SMA^+^ HSCs (n=21), and CD68^+^ Møs (n=37). A total of 115 ROIs were UV-illuminated. The released RNA probes were collected for PCR-amplification followed by AMPure XP beads (A63880; Beckman Coulter) purification. The purified libraries were sequenced by NextSeq 550 platform (Illumina) for spatial gene expression analysis.

### GeoMx data analysis

Fastq data were converted to DCC files by GeoMx NGS Pipeline software (Version 3.1.1.6) and analyzed by GeoMx DSP data center software (Version 3.1.0.222) for quality control and normalization. Quality control was assessed for each ROI to evaluate PCR contamination and sequencing depth. All 18,676 gene-associated probes and ROIs passed quality control. After Q3 normalization, gene expression data were analyzed by R (version 4.4.1) with the following packages: limma (version 3.60.4) for differential expression analysis, clusterProfiler (version 4.12.6) for gene set enrichment analysis, ggplot2 (version 3.5.1) for visualization, ComplexHeatmap (version 2.20.0) for heatmap generation and CellChat (version 1.6.1) for cell-cell communication analysis. Differential expression analysis between groups was conducted with |log2FoldChange| > 0.5 and Benjamini-Hochberg adjusted p-value < 0.05 as cutoff criteria. Adjusted p-values < 0.05 were considered statistically significant.

### Cells

HepaRG cells were obtained from Biopredic International (Saint Gregoire, France) and cultured in Williams’ E medium supplemented with 10% heat-inactivated fetal bovine serum (FBS), 5 µg/mL insulin, 50 µM hydrocortisone hemisuccinate, 1% L-glutamine, and 100 U/mL penicillin-streptomycin. BMOL cells were kindly provided by Dr. George Yeoh. Cells are maintained in Williams’ E medium supplemented with 10% FBS, 2mM L-glutamine, 100 U/mL penicillin/streptomycin, and 10 μg/mL Insulin (I0516, Sigma-Aldrich). Cells were seeded at a density of 1-5 × 10⁴ cells/cm² and cultured at 37 °C in a humidified incubator with 5% CO_2_, with the medium being changed every 48 to 72 hours.

### Cell counting kit-8 assay (CCK-8)

Cell viability was assessed by the Cell Counting Kit-8 (CK04-11, Dojindo Laboratories) according to the manufacturer’s instructions. Briefly, cells were seeded in 96-well plates at a density of 3-5×10^3^ cells per well and cultured overnight for attachment. After treatments under experimental conditions, 10μL CCK-8 solution was added to each well containing 100μL medium and incubated for 1-4 hours at 37 °C. The absorbance was measured at 450 nm by microplate reader (Spark 10M; TECAN).

### Colony formation assay

Colony formation ability was evaluated by colony formation assay. Cells were seeded in 6-well plates at 1000 cells per well and cultured for 2 weeks. Colonies were fixed with 4% paraformaldehyde and stained with 0.1% crystal violet for 20 minutes. Excess stain was gently washed off with distilled water, and then photographed.

### Cell cycle assay

Cells were synchronized by serum starvation overnight, followed by 24 h treatment under experimental conditions. After treatment, cells were harvested, fixed in ice-cold 75% ethanol overnight at -20°C, and stained with propidium iodide (50μg/ml) containing 100μg/ml RNase A for 1 hour at 37°C. Cell cycle distribution was analyzed using flow cytometry (FACSCanto II TPMA2, BD) with at least 10,000 cells recorded per sample. Data were analyzed by FlowJo software (version 10.8.1).

### RNA isolation and real-time PCR

Total RNA was isolated from cells using TRIzol reagent (15596018; Invitrogen) according to the manufacturer’s instructions, followed by chloroform phase separation (C2432; Sigma-Aldrich), isopropanol (1096342511; VWR) precipitation and 75% ethanol wash. The RNA pellet was dissolved in DEPC treated water (4387937; Thermo Fisher Scientific). RNA concentration was measured by multimode microplate reader (Spark 10M; TECAN). For cDNA synthesis, 1μg total RNA was reverse transcribed using RevertAid H Minus Reverse Transcriptase (EP0452; Thermo Fisher Scientific). Real-time PCR was performed on a StepOnePlus system (4376600; Thermo Fisher Scientific) by SYBR Green Master Mix (A25918; Thermo Fisher Scientific) with specific primers. The cycling conditions included initial denaturation at 95°C for 15 minutes, followed by 40 cycles of 95°C for 15 seconds, 60°C for 20 seconds, and 72°C for 20 seconds. Relative gene expression was calculated using the ΔΔCt method.

### Chromatin immunoprecipitation

Chromatin immunoprecipitation (ChIP) was performed by SimpleChIP® Enzymatic Chromatin IP Kit (#9003, Cell Signaling Technology) according to the manufacturer’s protocol. Briefly, cells were fixed with 1% formaldehyde for 10 minutes at room temperature. Chromatin was fragmented by partial digestion with Micrococcal Nuclease at 37°C for 20 minutes to obtain chromatin fragments of 1-5 nucleosomes. The chromatin was immunoprecipitated with c-Myc antibody (2 μg per sample) overnight at 4°C. After immunoprecipitation, protein G magnetic beads were added and incubated for 2 hours at 4°C. The beads were washed by low salt wash buffer three times and high salt wash buffer once. The protein-DNA complexes were eluted and cross-links were reversed by incubation with 5M NaCl at 65°C for 2 hours. After treatment with RNase A and Proteinase K, DNA was purified using spin columns and analyzed by qPCR.

### Immunohistochemistry

Paraffin-embedded tissue sections were deparaffinized by xylene (No. 9713.2, Carl Roth) and rehydrated by graded ethanol (K928.4; Carl Roth). For antigen retrieval, sections were heated in 10 mM citrate buffer (pH 6.0, C2404; Sigma-Aldrich) or 1 mM EDTA (pH 8.4, E9884; Sigma-Aldrich) by microwave (800W, alternating 30-second heating/cooling cycles for 15 minutes). After cooling to room temperature, sections were blocked with blocking peroxide (S200389-2; Dako) and 3% BSA (11930.03; SERVA Electrophoresis GmbH) in PBS (14190094; Thermo Fisher Scientific), then incubated with primary antibody in 0.3% BSA/PBS overnight at 4°C. After returning to room temperature, the sections were incubated with the secondary for 1 hour at room temperature. Immunoreactivity was visualized by 3,3’-diaminobenzidine (D5905; Sigma-Aldrich) with 0.03% hydrogen peroxide (1072102500; Merck), counterstained with hematoxylin (H-3401-500; Vector Laboratories). Finally, sections were dehydrated, mounted with mounting medium (C9368; Sigma-Aldrich), and scanned under microscope (DMRBE; Leica) or whole slide system (Aperio VERSA; Lecia). The staining intensity was quantified by measuring the integrated optical density per area (IOD/Area) by Image J software (Version 1.54d).

### Immunofluorescence staining

For immunofluorescence staining (IF), cells on coverslips were fixed with 4% formaldehyde solution (f1635; Sigma-Aldrich), permeabilized with 0.2% Triton X-100 (X100; Sigma-Aldrich), and blocked with 3% BSA in PBS. For tissue sections, samples were deparaffinized, rehydrated, and subjected to microwave-based antigen retrieval in 10 mM citrate buffer (pH 6.0) or 1 mM EDTA (pH 8.0) at 95-100°C (800W, alternating 30-second heating/cooling cycles for 15 minutes). Both cell and tissue samples were incubated with primary antibodies in 0.3% BSA/PBS overnight at 4°C, followed by secondary antibodies for 1 hour at room temperature. Sections were mounted using Fluoromount-G™ (00-4958-02; Thermo Fisher Scientific) and scanned with confocal microscope (TCS SP8; Leica).

### Western blotting

Total proteins were extracted by RIPA buffer containing protease inhibitor (04693132001; Roche) and phosphatase inhibitor cocktail 2 (P5726; Sigma-Aldrich), then quantified by BCA assay (5000116; Bio Rad). Twenty μg protein samples were separated by sodium dodecyl sulfate polyacrylamide electrophoresis at 80V through stacking gel and 120V through separating gel, then transferred to PVDF membranes (10600001; GE Healthcare Life Sciences) at 200mA for 2 hours. Following blocking with 5% BSA in Tris-buffered Saline with Tween (TBST), membranes were incubated with primary antibodies overnight at 4°C and HRP-conjugated secondary antibodies for 1 hour at room temperature. GAPDH were used as negative controls. Protein bands were visualized using Plus-ECL (NEL103001EA; Perkin Elmer) and detected by the imaging system (Fusion SL4; PEQLAB).

### Statistical analysis

All data were expressed as mean ± standard deviation (SD). Statistical analyses were performed using Student’s t-test for comparisons between two groups and one-way ANOVA for multiple group comparisons. Statistical significance was defined as P < 0.05, with significance levels indicated as follows: *, P < 0.05; **, P < 0.01; ***, P < 0.001 and ****, P < 0.0001.

The detailed information on materials and reagents is summarized in **Table S1**.

## Acknowledgements

We are grateful to Drs. Bin Gao and Dechun Feng to provide support in animal experiments and constructive discussion and Dr. George Yeoh for kindly providing BMOL cells. We thank the Human Tissue and Cell Research Foundation, a nonprofit foundation regulated by German civil law, which facilitates research with human tissue through the provision of an ethical and legal framework for prospective sample collection. We acknowledge the support of the LIMA Live Cell Imaging at Microscopy Core Facility Platform Mannheim (CFPM).

## Author’s contributions

H.L.W. conceived and designed the project. H.L., C.S., and H.D. collected the patient samples and performed pathological evaluation. C.T. and T.L. undertook experiments. C.T., and H.W. performed bioinformatic analyses. C.T. and H.L.W. drafted the article. C.T., S.H., R.L., M.P.A.E., H.D., S.D., and H.L.W. discussed the data and edited the article critically.

## Supplementary data

**Figure S1.**
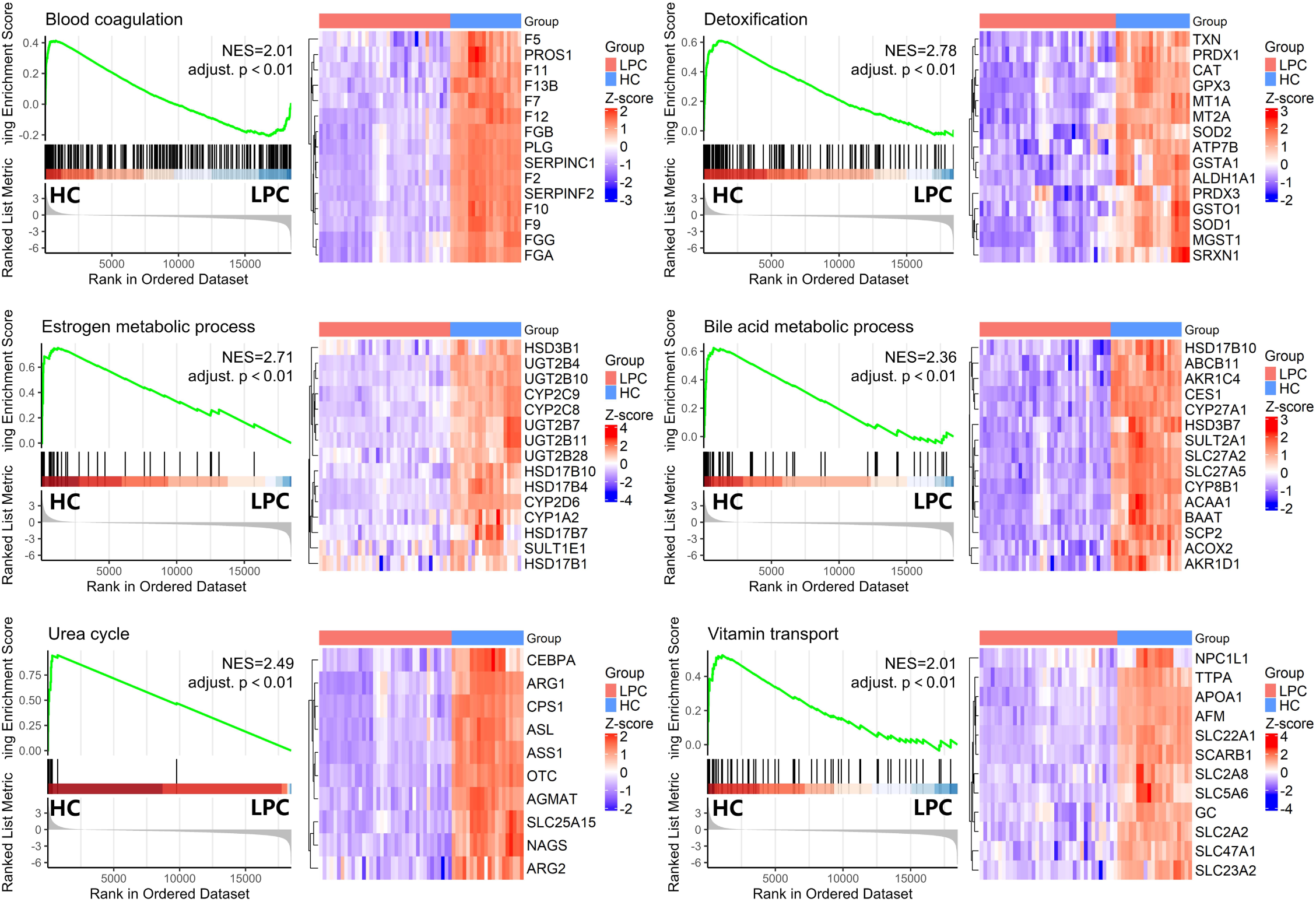
GSEA identifies downregulation of hepatic metabolic and functional pathways in LPCs compared to HCs. GSEA was performed to compare the transcriptomic profiles of LPCs and HCs. Enrichment plots (left panels) illustrate curated gene sets related to liver-specific functions. Normalized enrichment scores (NES) and adjusted *p* values are provided for each comparison. The right panels display heatmaps of representative genes within each pathway, with z-score normalization applied across samples.

**Figure S2.**
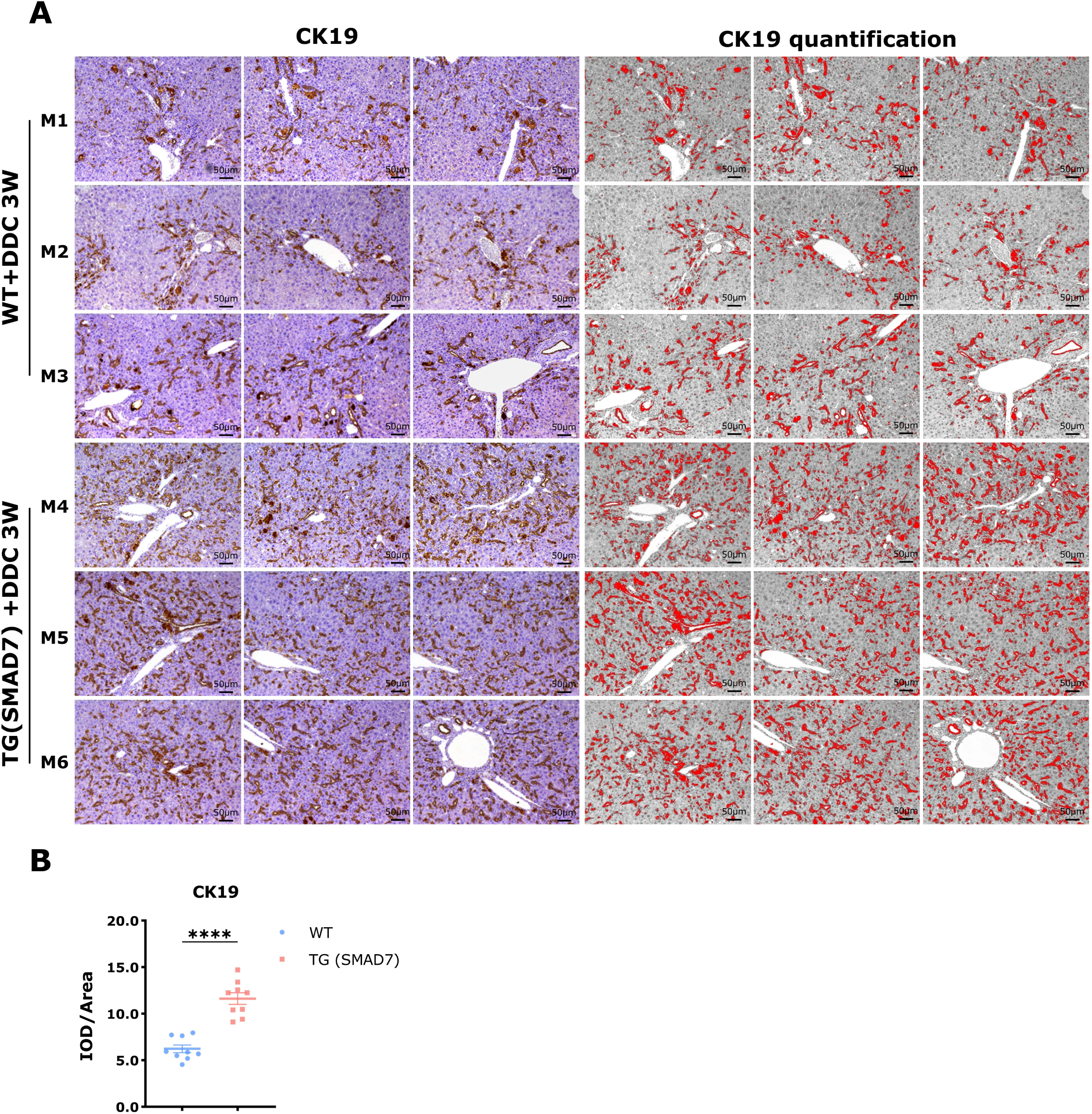
Overexpression of SMAD7 increases DDC-induced LPCs proliferation. **(A)** IHC staining was performed on liver tissues from wild-type and SMAD7 transgenic mice fed with DDC for 4 weeks. Representative images depict CK19 expression in liver sections from three individual wild-type mice (M1-M3) and three SMAD7 transgenic mice (M4-M6). For each mouse, three different areas were analyzed. The left panels show the original CK19 staining, while the right panels display digital quantification of CK19-positive areas (highlighted in red) using ImageJ software. Scale bars: 50 μm. **(B)** Quantification of CK19 expression was performed using integrated optical density (IOD) normalized to tissue area. Data are shown as mean ± SD. *p < 0.0001 (****)*.

**Figure S3.**
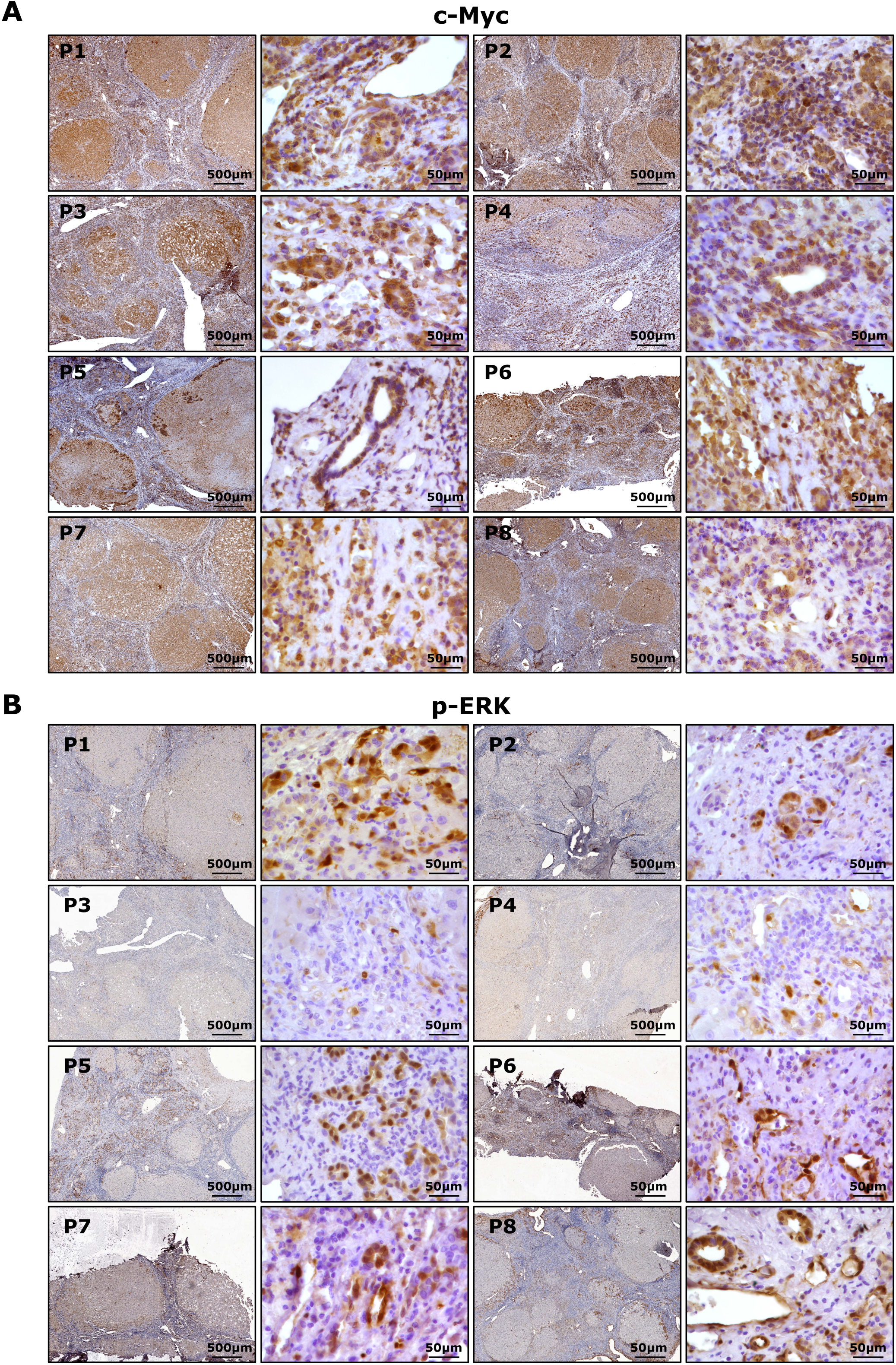
Activation of c-Myc and p-ERK in ALF patients. (A-B) IHC staining for c-Myc and p-ERK was performed on liver tissues from eight ALF patients. Each patient sample includes an overview image and areas with higher magnification. Scale bars: 50 μm and 200 μm.

**Figure S4.**
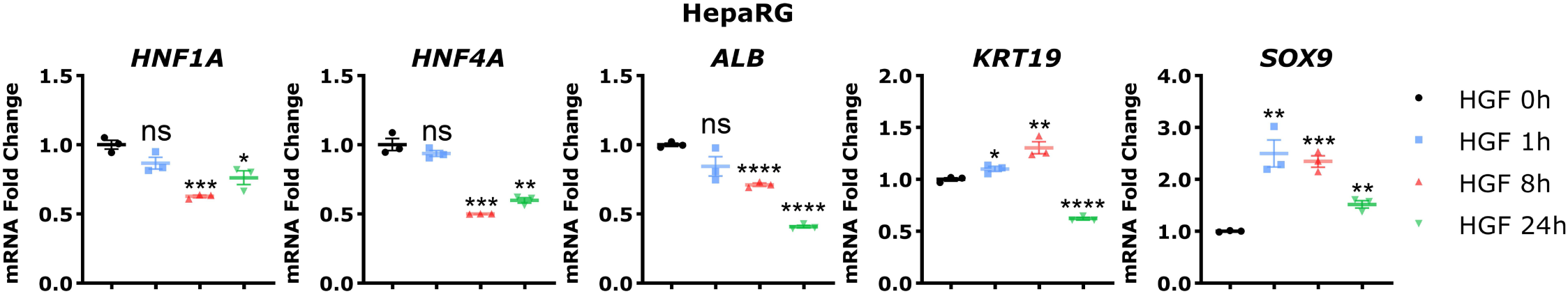
The effects of HGF on the expression of hepatocyte and cholangiocyte genes in HepaRG cells. qRT-PCR was used to measure mRNA expression of Hnf1a, Hnf4a, and Sox9 in HepaRG cells treated with 20 ng/mL HGF for 0, 4, 8, 12 and 24 h. Data are shown as mean ± SD. *p < 0.05 (*), p < 0.01 (**), p < 0.001 (***), p < 0.0001 (****)*; ns, not significant.

**Table S1.**
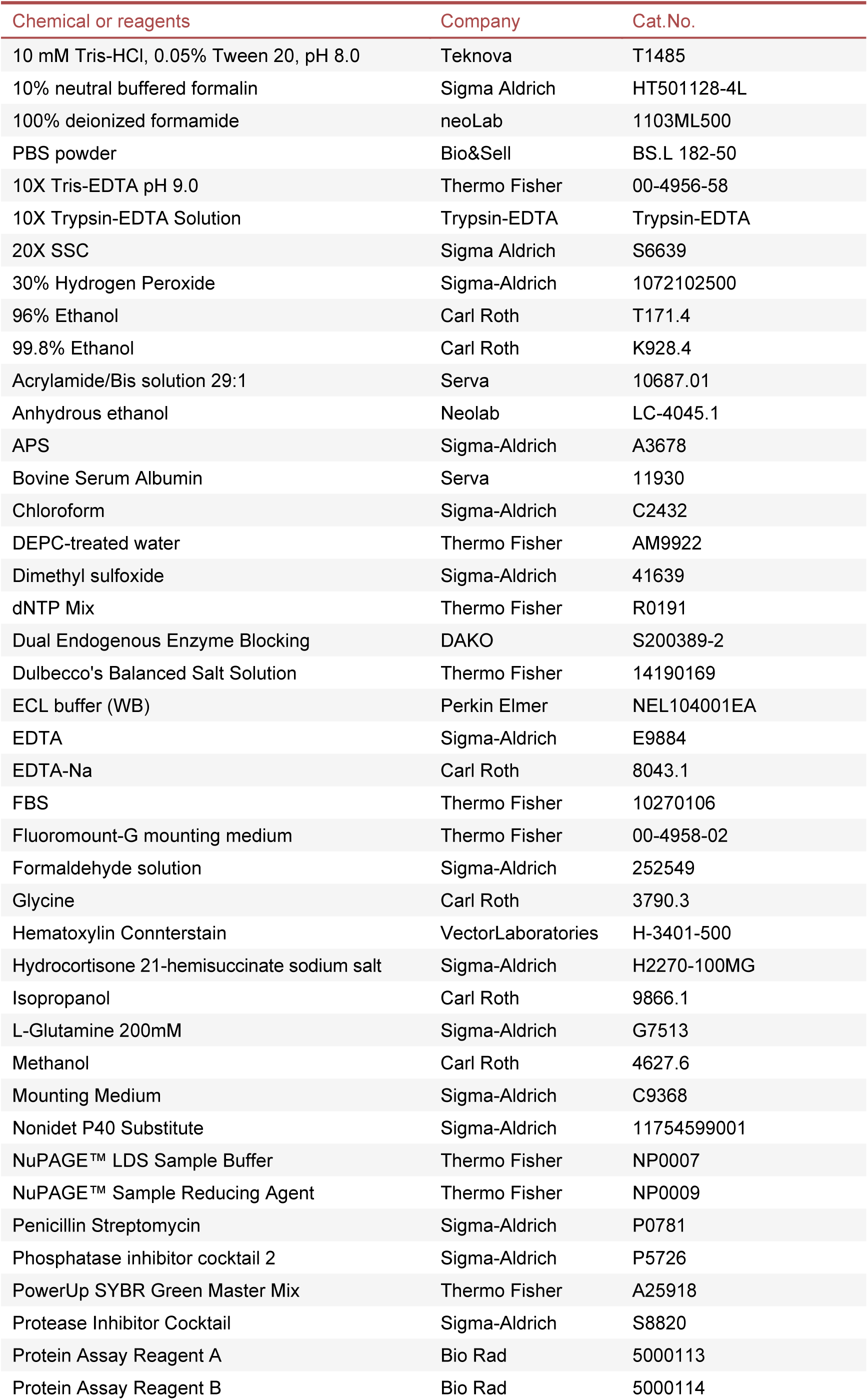
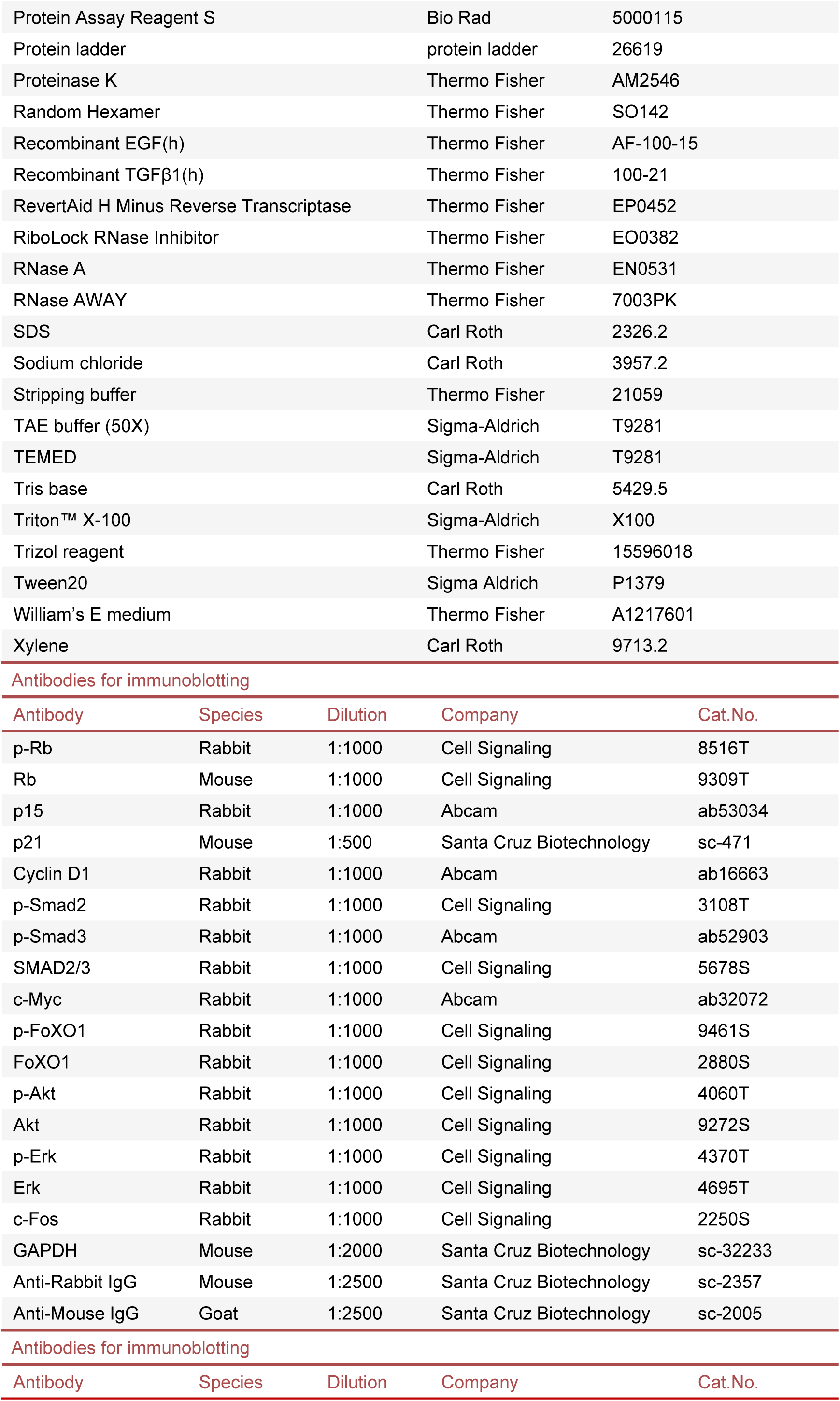
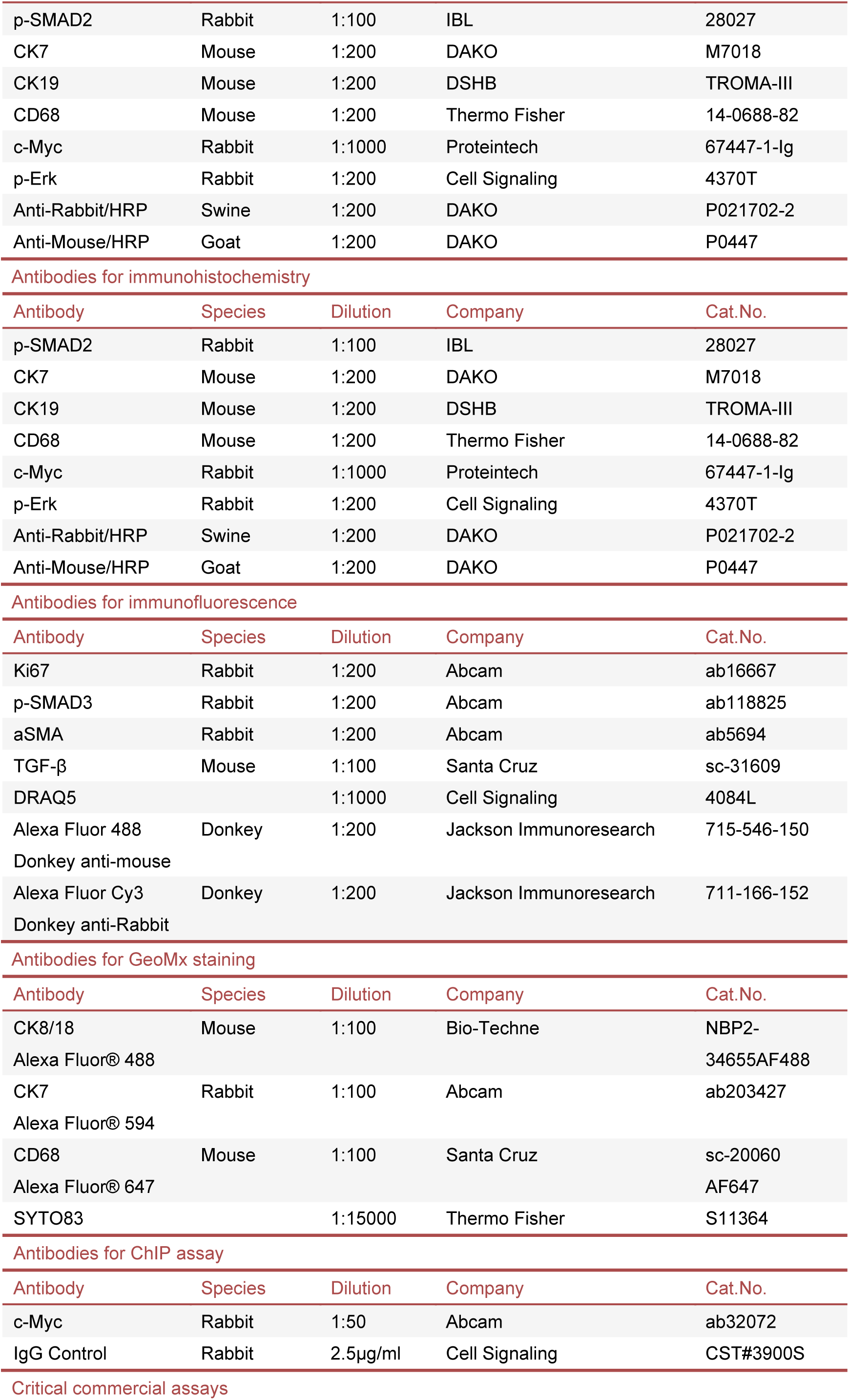
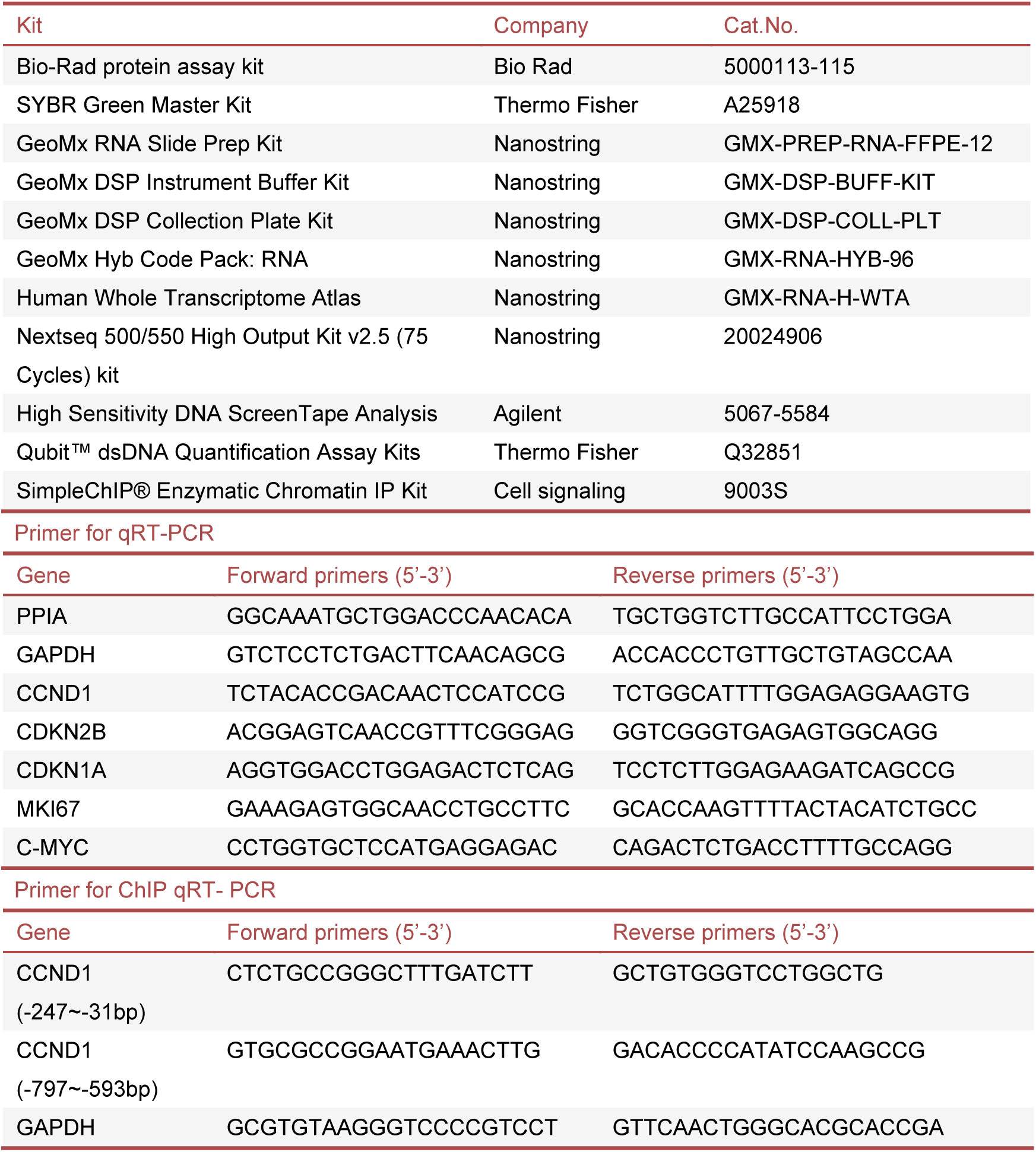
Materials, reagents, antibodies, commercial kits, and primers.

